# Discovery of an unusual high number of *de novo* mutations in sperm of older men using duplex sequencing

**DOI:** 10.1101/2021.04.26.441422

**Authors:** Renato Salazar, Barbara Arbeithuber, Maja Ivankovic, Monika Heinzl, Sofia Moura, Ingrid Hartl, Theresa Mair, Angelika Lahnsteiner, Thomas Ebner, Omar Shebl, Johannes Pröll, Irene Tiemann-Boege

## Abstract

*De novo* mutations (DNMs) are an important player in heritable diseases and evolution. Of particular interest are highly recurrent DNMs associated with congenital disorders that have been described as selfish mutations expanding in the male germline, thus becoming more frequent with age. Here, we have adapted duplex sequencing (DS), an ultra-deep sequencing method that renders sequence information on both DNA strands; thus, one mutation can be reliably called in millions of sequenced bases. With DS, we examined ∼4.5 kb of the *FGFR3* coding region in sperm DNA from older and younger donors. We identified sites with variant frequencies of 10^−4^ to 10^−5^, with an overall mutation frequency of the region of ∼6×10^−7^. Some of the substitutions were re-current and were found at a higher variant frequency in older donors than in younger ones, or exclusively, in older donors. Also, older donors harbored more mutations associated with congenital disorders. Other mutations were present in both age groups suggesting that these might result from a different mechanism (e.g., post-zygotic mosaicism). We also observed that independent of age, the frequency and deleteriousness of the mutational spectra was more similar to COSMIC than to gnomAD variants. Our approach is an important strategy to identify mutations that could be associated with a gain-of-function of the receptor tyrosine kinase activity, with unexplored consequences in a society with delayed fatherhood.

## Introduction

There are certain *de novo* mutations (DNMs) that are highly recurrent with mutation rates orders of magnitude higher than genome average. These mutations have been discovered because of their association with congenital disorders. Moreover, these mutations have several other associated characteristics (reviewed in (Arnheim and Calabrese 2009; Goriely et al. 2009; Goriely and Wilkie 2012; Arnheim and Calabrese 2016)): they encode missense substitutions with gain-of-function properties (activating mutations); they occur exclusively in the male germline; and older men have a higher probability of having an affected child than younger males, known as the paternal age-effect (PAE) described already decades ago (Risch et al. 1987; Crow 2000; Crow 2012). In the literature, these mutations have been termed as PAE-mutations, selfish, or RAMP (recurrent, <u>a</u>utosomal dominant, <u>m</u>ale biased, and <u>p</u>aternal age effect) mutations, as reviewed in (Arnheim and Calabrese 2009; Goriely and Wilkie 2012; Arnheim and Calabrese 2016).

Over the last decades, it was shown that the high incidence levels and steep increase with age of these PAE-mutations are not solely due to errors occurring during the continuous replicative process of spermatogenesis (Risch et al. 1987; Crow 2000; Crow 2012). Instead, it was suggested that these behave like driver mutations, well known in cancer, which are mutations that promote their own clonal expansion. In particular, PAE-mutations have been described in genes such as *FGFR2, FGFR3, HRAS, PTPN11, KRAS* and *RET* (Qin et al. 2007; Choi et al. 2012; Shinde et al. 2013; Maher et al. 2016), and more recently in six new genes (*BRAF, CBL, MAPK1, MAPK2, RAF1*, and *SOS1*) (Maher et al. 2018), all acting in the receptor tyrosine kinase (RTK)-RAS signaling pathway and expressed in spermatogonial stem cells (SSCs). The mutations modify the signal modulation of the RTK-RAS pathway by an activating effect of the mutant protein. This dysregulation of the RTK-RAS pathway drives the preferential expansion of mutant SSCs, observed in the testis as mutant clusters that become larger with age (Qin et al. 2007; Choi et al. 2008; Choi et al. 2012; Shinde et al. 2013; Yoon et al. 2013; Maher et al. 2016; Maher et al. 2018). Therefore, as men age, the germline becomes a mosaic for multiple PAE-mutations, all in different anatomical locations of the testes (Shinde et al. 2013; Arnheim and Calabrese 2016; Maher et al. 2018), suggesting that each mutation arises and expands independently.

The clonal expansion of some PAE-mutations in the testis with age also results in more mutant sperm in older individuals, and an increased incidence of the associated congenital disorder with paternal age, as observed for example for achondroplasia (ACH), a growth disorder with a defect in FGFR3 signaling (Risch et al. 1987; Tiemann-Boege et al. 2002; Shinde et al. 2013). Interestingly, the transmission of mutations in SSCs to sperm and hence to the offspring does not always correlate positively, especially mutations associated with tumors hardly overlap with spontaneous germline mutations (Goriely et al. 2009; Aoki and Matsubara 2013; Giannoulatou et al. 2013).

In spite of the importance of PAE-mutations in the male germline which is highlighted by their high incidence, increased frequency with paternal age, and phenotypes potentially leading to early or late-onset disorders in children of older men, we know very little about this mutagenic mechanism. There are still many open questions on how and to what extent specific PAE-mutations affect cell growth and spermatogonial stem cell differentiation, and the role of apoptosis or cell-death counterbalancing clonally expanding cells. So far, studies have focused on well-characterized mutations associated with a disorder (reviewed in (Arnheim and Calabrese 2009; Goriely et al. 2009; Goriely and Wilkie 2012; Arnheim and Calabrese 2016)) or on the most prevalent mutations observed at late oncogenic stages in somatic tumors, which were shown to be unrelated to the selfish expansion of mutant spermatogonial stem cells (Goriely et al. 2009; Giannoulatou et al. 2013; Maher et al. 2016). However, many genes in the RTK-RAS signaling pathway, or in other pathways, could harbor as yet unknown mutations that could be expanding with paternal age but have gone undetected so far, though, with important health consequences in children of older fathers. To overcome this, a more recent study used a sequencing strategy to discover prospective driver mutations in genes of the RTK-RAS pathway by enriching for mutant SSCs identified in testes biopsies by a high RTK activity that identified 61 mutations at a frequency of 0.06% or larger (Maher et al. 2018).

In this work, we further developed the discovery of prospective RTK mutations in the male germline to include ultra-low frequency variants (with a variant frequency of <2 × 10^−5^). For this purpose, we used sperm since this biological material is easy to sample and provides information on whether mutations are passed on throughout the different maturation and differentiated stages of the male germline. Accurately identifying ultra-low frequency mutations by sequencing is still technically very challenging, especially if no enrichment of mutants is possible (e.g., high RTK signaling). To date, only a few methods can achieve the required sensitivity and accuracy. One of these methods is duplex sequencing (DS), a strategy that organizes sequence reads derived from the same DNA molecule into families with information on the forward and reverse strands (Schmitt et al. 2012). Given the excessive sequencing depth required in DS, in this work we focused on the coding region of *FGFR3* because (1) this gene has well characterized PAE-mutations; (2) various reported DNMs associated with congenital disorders have incidence levels that fall within the sensitivity of DS (reviewed in (Goriely et al. 2009; Goriely and Wilkie 2012; Shinde et al. 2013; Maher et al. 2018)); and (3) *FGFR3* has been categorized as an oncogene with many missense mutations and high oncogenic score (Tomasetti and Vogelstein 2015).

We sequenced the coding regions of the *FGFR3* gene at an average coverage depth of ∼17,000x (up to ∼38,000x) in young and an old donor pools. We identified 75 unique variants at 72 different genomic positions in the exonic *FGFR3* sequence (exon 3-15 without exon 11); of those, 20 have been reported in other databases to be associated with congenital disorders (Human Gene Mutation database (HGMD)) and/or tumors (Catalogue Of Somatic Mutations In Cancer (COSMIC)); 34 DNMs have never been reported before (Table S1). Some DNMs were found exclusively or at a higher frequency in the older donor set. Interestingly, we also identified many higher frequency variants independent of the age of the donors, suggesting that these might not be testis-exclusive mutations, but could have potentially happened earlier in development from a post-zygotic event. Overall, our method is a powerful approach to identify PAE-mutations that can be then further characterized to better understand this important type of mutagenesis in the male germline.

## Results

### DS detects ultra-low mutation frequencies

Short-read next-generation sequencing (NGS) is a widely used method for prospective screening of mutations, but given its high error rate of ∼0.1-2%, it cannot be used for detecting low or ultra-low frequency variants or mutations (reviewed in (Salk et al. 2018)). In the past years, different approaches for library preparation have been published to distinguish technical errors from real DNA variants, decreasing error rates to ∼10^−5^ (depending on the method) (Ebner et al. 2011; Jabara et al. 2011; O’Roak et al. 2012; Schmitt et al. 2012; Hiatt et al. 2013; Lou et al. 2013). This is achieved by individually tagging each DNA molecule before amplification and sequencing with an Illumina platform. The tags are then used for grouping reads into a consensus sequence (family). A true mutation is present in the majority of the reads within a family, whereas PCR and sequencing errors are only present in a subset of reads.

The sequencing method with the lowest reported error rate using such a tagging approach is DS, which uses a double barcode strategy to retrieve information from both strands of the original DNA molecule. In DS, reads are first organized into families of either the forward or the reverse strand, known as single-stranded consensus sequence (SSCS). Then, the complement SSCS are further grouped into the duplex consensus sequence (DCS) representing both DNA strands of one initial DNA molecule (Figure 1A). A variant is considered true, if present in the majority of reads of both strands in the DCS, allowing for an extremely low error rate of ∼10^−7^ to 10^−8^ which is a major improvement of at least 1-2 orders of magnitude compared to other ultra-deep sequencing methods (Schmitt et al. 2012; Fox et al. 2014; Kennedy et al. 2014). Most ultra-deep sequencing methods use information of only one strand and are limited mainly by DNA lesions (Lou et al. 2013; Arbeithuber et al. 2016).

**Figure 1.**
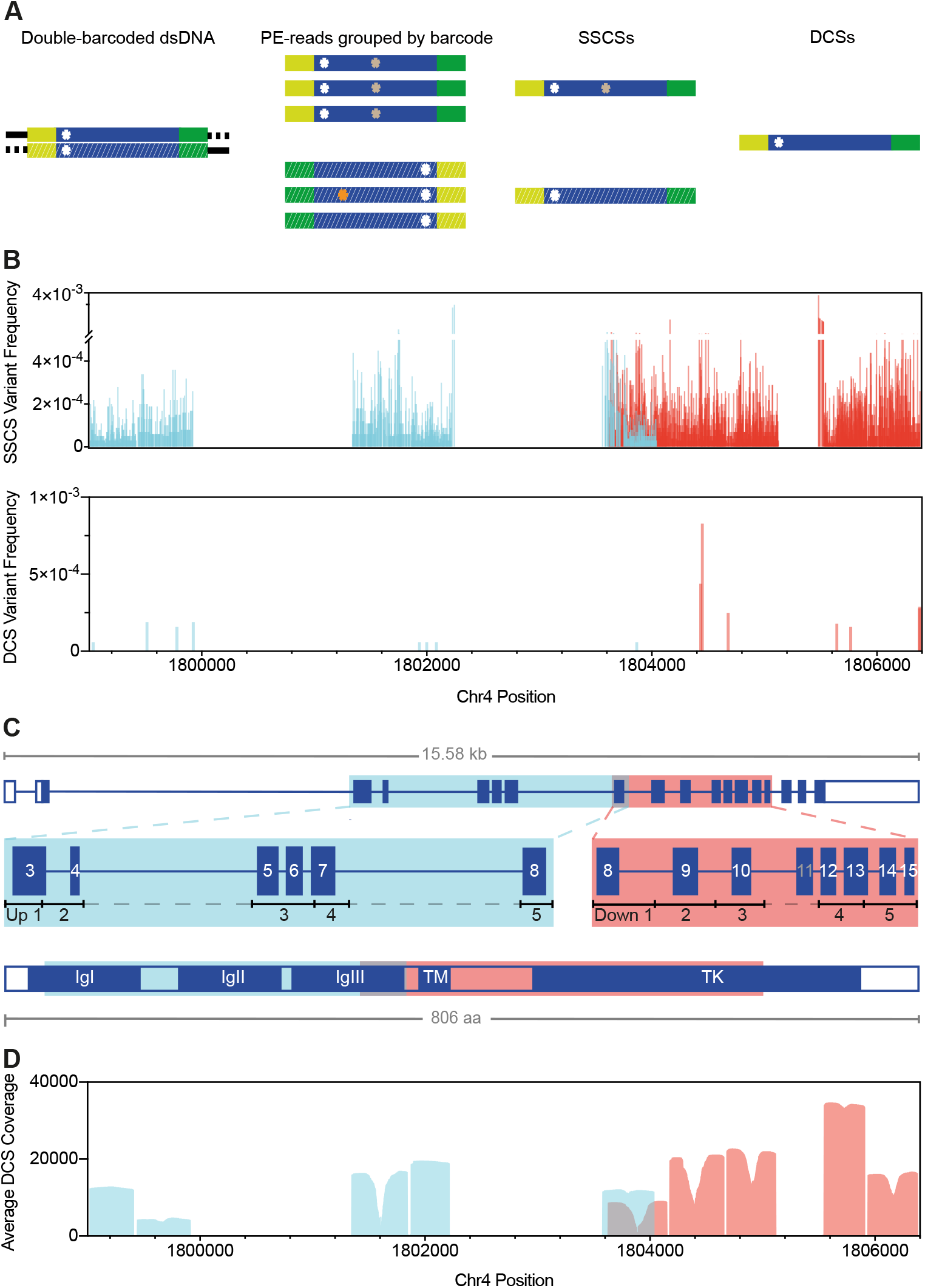
DS using *FGFR3* targeted sequencing strategy with enzymatic digestion. (*A*) DS overview: barcoded adapters (yellow and green) were ligated at both ends of size-selected restriction enzyme digested genomic fragments (blue). After several rounds of PCR amplification and hybridization captures, libraries were sequenced on the Illumina MiSeq (v3 600-cycles); for sequence analysis, paired-end (PE)-reads were grouped into Single Strand Consensus Sequences (SSCS) and complementary SSCS were joined into a Duplex Consensus Sequence (DCS). Only DCS with mutations in both complementary SSCS (white asterisk) were considered true variants. Colored asterisks represent artefacts (e.g., sequencing and PCR errors). (B) Variants detected at 2,881 different positions of the total 4,405 sequenced positions identified in SSCS (n=14,552 total variants; VAF = 1.0 × 10^−5^ to 3.8 × 10^−3^) or at 15 positions in DCS (n=15 total variants; VAF=6.0 × 10^−5^ to 8.3 × 10^−4^) of two *FGFR3* libraries (FGFR3 Up O Oct19 Re-seq and FGFR3 Down O BAT), each targeting either the Up 1-5 (blue) or the Down 1-5 (red) regions/subregions. (*C*) Exonic structure of *FGFR3*. Shown are the five upstream (blue) or downstream (red) regions targeted via restriction enzyme fragmentation that include exons 3 to 8 or 8 to 15 (except exon 11), respectively; restriction digests rendered fragments ∼370-550 bp in size. Acronyms of the FGFR3 structure: Immunoglobulin-like domain I-III (IgI-III); Transmembrane domain (TM); Tyrosine kinase domain (TK); amino acids (aa). (*D*) Average DCS coverage per position of all sequenced libraries.

These DNA lesions can be amplified during the library preparation resulting in false positives. DS greatly reduces errors resulting from DNA damage since a true variant is present in both DNA strands, while a DNA lesion is only found in one DNA strand (Arbeithuber et al. 2016).

Here, we illustrate the higher accuracy of a duplex over a single-stranded consensus sequence (DCS vs. SSCS), the latter being equivalent to ultra-deep sequencing approaches examining only one DNA strand. We compared the mutation frequency obtained when grouping reads into a family from either the forward or the reverse strand (SSCS) with the mutations obtained from the DCS (Figure 1B). In SSCS, we observed many more variants, also with a higher variant allele frequency (VAF) that were filtered out in the DCS (Figure 1B). Note that insertion-deletions were not considered here. Some variants within SSCS could be PCR jackpots (errors occurring during the initial PCR cycles that are exponentially amplified (Lou et al. 2013)) or alternatively the product of DNA lesions as observed previously (Arbeithuber et al. 2016). In fact, the majority (86%) of the called SSCS were C>A transversions (the product of oxo-G lesions) or G>A / C>T transitions (the product of cytosine deamination) as shown in Supplemental Figure S1.

The high accuracy of DS is tied with the disadvantage that this approach is very data expensive and only a fraction of the input molecules ends up in a DCS. For this reason, we only focused on exon 3 to 15 of the *FGFR3* gene (∼4.5 kb). As shown in Figure 1C, each of our sequencing libraries targeted one of two ∼2.5 kb regions of *FGFR3*. The two regions were divided further into 5 sub-regions (Up 1-5 or Down 1-5) each ∼500 bp in size that could be fully covered by the two paired-end (PE)-reads with an Illumina MiSeq v3 600-cycles sequencing kit. Our DS strategy also included a new approach that enriched for these sub-regions using restriction enzymes followed by the removal of bulk DNA via bead size selection (see Methods). With this DS strategy, we achieved a median DCS coverage of ∼10,692 (median DCS coverage per library ranges from 6,222 to 38,422); see Figure 1D and Supplemental Table S1).

In order to confirm that variants called by our pipeline represent true mutations, six of the mutations identified with DS were measured with other ultra-sensitive detection methods, namely bead emulsion amplification (BEA) as described in (Shinde et al. 2013) or droplet digital PCR (ddPCR); see Supplemental Table S2. BEA is an in-house digital PCR method based on the amplification of single molecules on magnetic beads within an emulsion (Boulanger et al. 2012; Shinde et al. 2013). The ddPCR technology (Bio-Rad Laboratories) is based on the partitioning of PCR reactions into thousands of individual reactions within water-in-oil droplets that are read out in an automated droplet flow-cytometer after the PCR reaction is concluded (Hindson et al. 2011); see Methods for further information. Our DS measurements were similar among methods; albeit, given the low number of events captured in DS the confidence intervals are larger for DS than for ddPCR. Moreover, in BEA and ddPCR more starting molecules were used (∼300,000 genomes) making these two methods more accurate for the quantification of very low mutation frequencies compared to DS. A Fischer’s exact test for all possible comparisons did not reveal any significant difference between the measured mutation frequencies.

As a further demonstration of the high sensitivity and accuracy of DS, we carried out a mixing experiment. For this purpose, different genomic DNA obtained from Coriell Cell Repositories (Camden, NJ) each carrying a point mutation (c.742C>T, c.746C>G, c.749C>G, and c.1620C>A) were spiked into DNA from a younger donor at ratios from 1/10 to 1/10,000 in steps of one order of magnitude (Figure 2). With this orthogonal assay, we demonstrated that the variant frequency detected with DS was equivalent to the expected input of the 10-fold dilutions. Note that the starting input amounts varied due to pipetting errors and the occasional inaccuracy of spectrophotometric measurements used to establish the initial DNA concentration. Fourteen out of 16 variants (four variants per library) were identified. Two replicates of the 1:10,000 dilutions were not detected likely due to sampling drop-out at the Poisson distribution level. The Pearson’s correlation coefficient between observed and expected input dilutions of the 14 measurements was very high (R^2^ = 0.96, p-value = 9.8 × 10^−10^) demonstrating the power of DS to measure ultra-rare variants (Figure 2).

**Figure 2.**
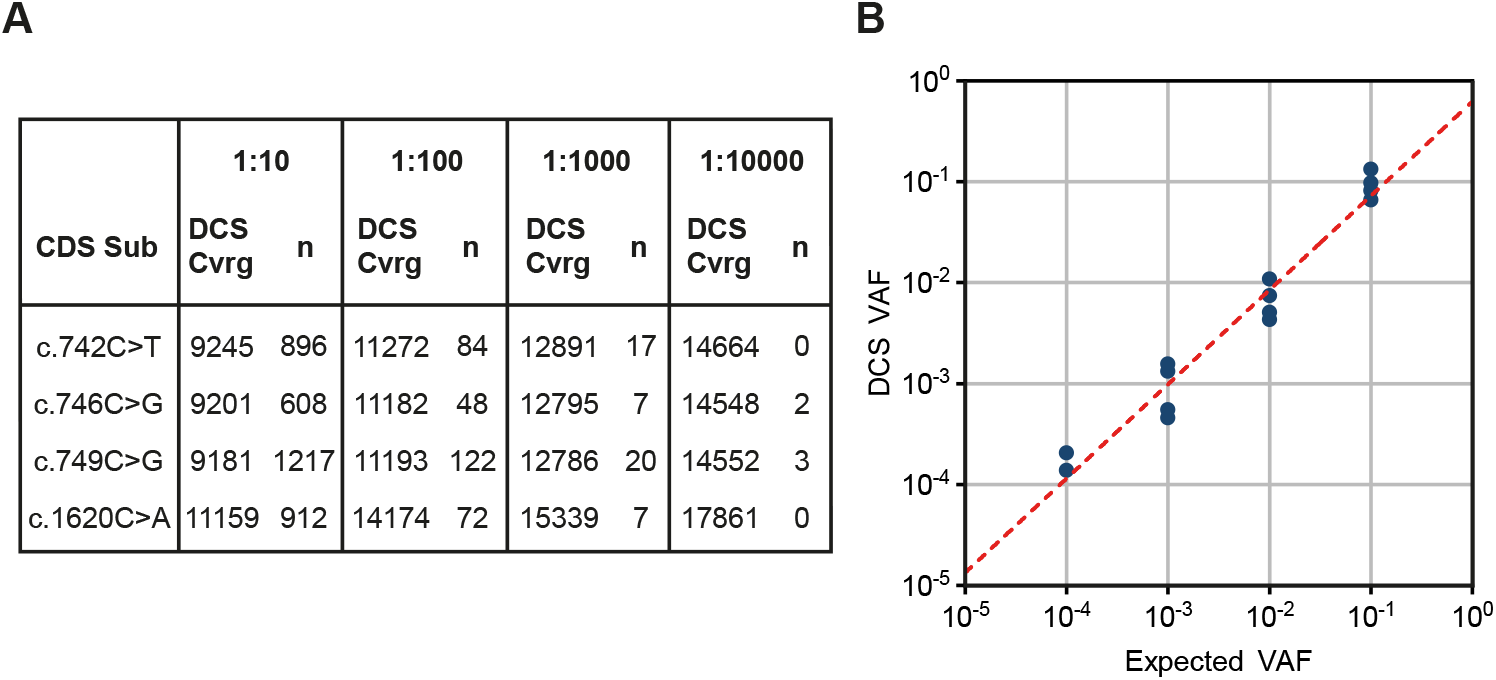
DS spike-in test. (*A*) DNA from 4 different cell lines (Coriell Cell Repositories; Ref: CD00002, NA00711, NA08909, GM18666) was serially diluted into sperm DNA from a young donor in order to prepare four DS libraries (“FGFR3 Y C 1:10”, “FGFR3 Y C 1:100”, “FGFR3 Y C 1:1000”, “FGFR3 Y C 1:10000”) with expected mutant frequencies of 1:10, 1:100, 1:1000, and 1:10000, respectively. DCS coverage and the number of times the variants were found in each library are shown. (*B*) Expected VAF vs. measured VAF with DS; 14 out of the 16 mutations were detected; R^2^ = 0.96 (Pearson’s correlation coefficient).

### *Substitutions detected in the coding region of* FGFR3 *in sperm DNA*

In total, we sequenced 12 libraries, each prepared from sperm DNA pooled from five healthy individuals, mainly of European ancestry. Six libraries featured younger donors with ages ranging from 26 to 30 years, and the remaining six featured older donors with ages ranging from 49 to 59 years; see Supplemental Table S3 and Supplemental Table S4. The distribution of donor ages in the young and old pool was significantly different (Supplemental Figure S2). Further, in spite of differences in the library preparation protocols, we did not observe differences in the overall mutation frequency among libraries, suggesting that our approach is quite robust (Supplemental Figure S3).

Consensus sequences were first formed using Du Novo (Stoler et al. 2016; Stoler et al. 2020) and variants were called using the Naive Variant Caller (Blankenberg et al. 2014), followed by a more thorough screening with our Variant Analyzer software (Povysil et al. 2021). The Variant Analyzer re-examines raw reads from Du Novo variant calls and provides different summary data that categorizes the confidence level of a variant by a tier-based system. This tier classification is based mainly on the number of reads per family and the proportion of the alternative allele in the consensus sequence (e.g., Tier 1 is composed by families with 3 or more members for both forward and/or reverse strands). With this tier classification (see Supplemental Table S5), we can distinguish true mutants from bad quality data or biases leading likely to false positives. Moreover, our Variant Analyzer supports the analysis of reads without a family as demonstrated in (Povysil et al. 2021), which allowed maximizing the data output.

In total, we identified 395 mutations classified as Tier 1 or Tier 2 located at 380 different genomic positions of the 4,405 sequenced positions of *FGFR3*. This list does not include single-nucleotide polymorphisms (SNPs) (variants with a frequency of ∼10% in the donor pool, likely to be a heterozygous position in one of the individuals) or variants with DCS coverage below 1000. We shortlisted the 395 mutations to those located in exons. We only included Tier 2 variants when detected at least twice with a Tier 1 and/or Tier 2 in the same or different libraries. Multiple hits increase the likelihood for these to be real. We retained 99 high-confidence variant counts (75 unique) at 72 genomic positions (Figure 3 and Supplemental Table S1).

**Figure 3.**
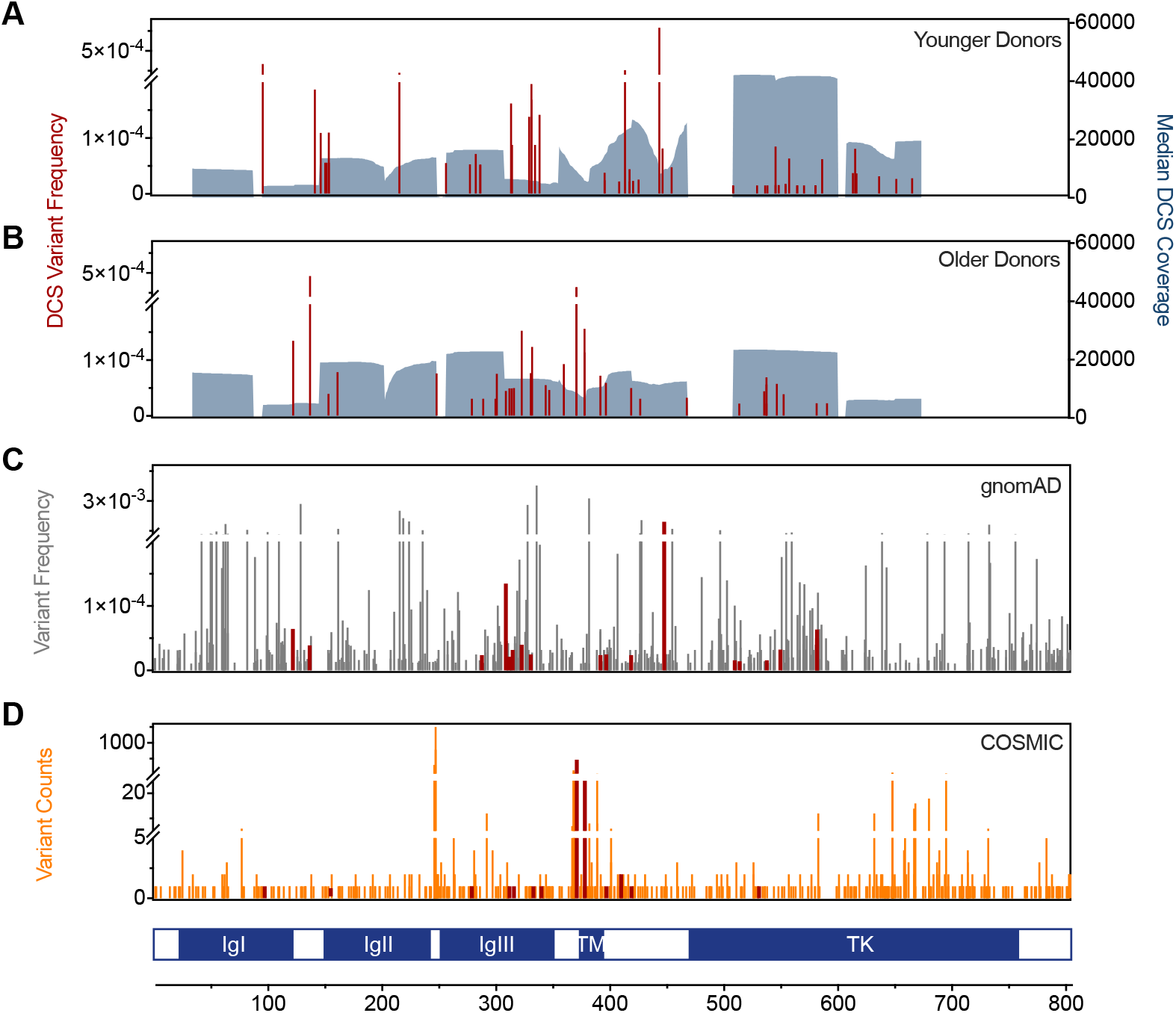
Variant frequency and DCS coverage. In red, the variant allele frequency (VAF) of the 99 exonic variant counts found in all libraries from younger (58 counts, (*A*)) and older donors (41 counts, (*B*)) listed in detail in Supplemental Table S1. DCS coverage (grey shaded area) is the median DCS coverage at exonic positions of all libraries within the same age group. (*C*) Exonic gnomAD variants for *FGFR3* with frequencies between 1 × 10^−5^ and 1 × 10^−2^. (*D*) Counts of exonic variants associated with tumors (orange) were retrieved from the COSMIC database. The gnomAD and COSMIC variants detected also by DS are shown in red. Immunoglobulin-like domain I-III (IgI-III); Transmembrane domain (TM); Tyrosine kinase domain (TK) of FGFR3.

The mutants detected in this study had a VAF (estimated as the ratio of the alternate allele divided by the reference allele or mutation count per depth of coverage at a given genomic position) between 1.5 × 10^−5^ to 7.9 × 10^−4^ (Figure 3 and Supplemental Table S1). Of the 75 discovered variants, 34 of them have not been reported before in any public databases, 13 were unique substitutions associated with tumors (mainly bladder, skin, and multiple myeloma, which are the most common cancer associations within *FGFR3*), and 32 were substitutions reported in the gnomAD database (likely viable mutants).

We also calculated the overall mutation frequency, which is an estimate of the total number of variants per number of sequenced base pairs, and can be used to compare overall mutation frequencies with other methods, or differences between donor groups or domains (Figure 4). Note that this mutation frequency or substitution/base pair represents *de novo* mutation (DNM) events, and was estimated as described in Methods. As shown in Figure 4A, the mutation frequency of the exonic region of *FGFR3* is ∼4.58 × 10^−7^, which is one order of magnitude higher than measured genome-wide with DS (1 × 10^−8^) ((Abascal et al. 2021)) or by sequencing trio pedigrees (1.2 × 10^−8^) ((Kong et al. 2012)). Unlike genome-wide data, *FGFR3* had a particularly higher frequency of transversions and non-CpG transitions (Supplemental Figure S4), suggesting a unique mutagenic mechanism in this gene explored in more detail in the next sections. Interestingly, the number of missense mutations was significantly higher compared to synonymous, stop-gained or splice region variants with approximately two-thirds of the missense variants being deleterious based on the SIFT score (Supplemental Figure S5). Also, deleterious mutations or mutations with unknown effect based on the SIFT score were overall more frequent than tolerated/benign variants (Figure 4A).

**Figure 4.**
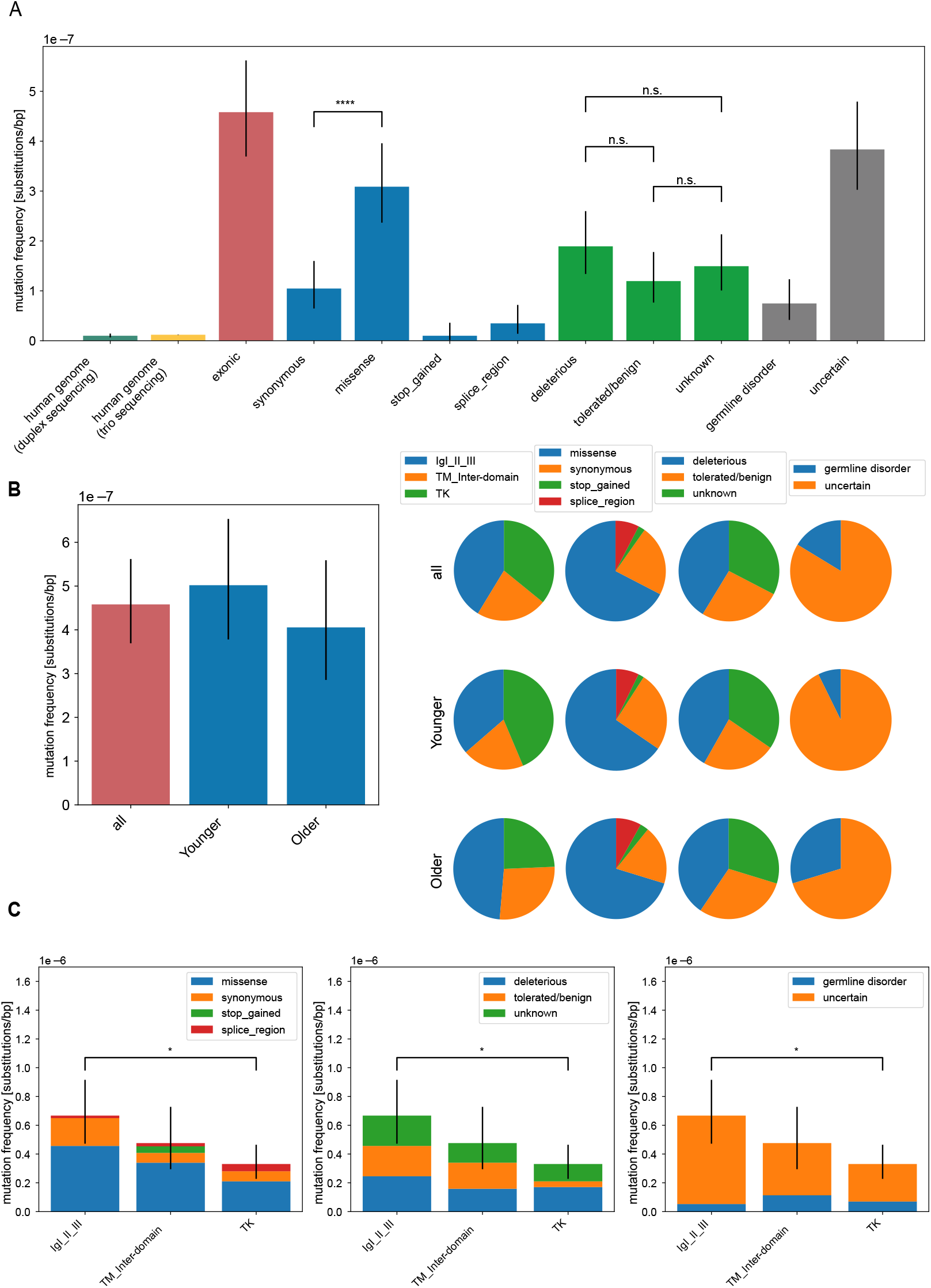
Analysis of mutation frequencies in *FGFR3*. Overall mutation frequencies estimated as number of variants (considered as DNM) divided by the number of sequenced nucleotides of the targeted regions ***(A*) Mutation frequencies (substitutions per base pair)** measured in the human genome by duplex sequencing (Abascal et al. 2021) and trio sequencing ((Kong et al. 2012)) and compared to the exonic regions of *FGFR3* (n=92). We further subdivided the mutations in the type of amino acid substitution (<u>synonymous n=21, missense n=62</u>, <u>stop_gained n=2, splice_region n=7)</u>, the effect on the protein based on the SIFT score (deleterious n=38, tolerated/benign n=24, unknown n=30) or with a phenotype according to the HGMD database (germline disorder n=15, uncertain n=77) listed in detail in Supplemental Table S1. **(*B*) Age dependency**. Mutation frequencies compared between donor groups (all n=92, Younger n=55, Older n=37). The pie chart is based on the mutation frequency for each group. **(*C*) Domain analysis**. Mutation frequencies compared among protein domains (IgI-III n=38, TM_Inter-domain n=21, TK n=33) categorized after substitution type, deleteriousness, and associated germline disorder. Error bars are confidence intervals of a Poisson distribution. For pairwise testing the Chi-square test with Bonferroni-Holm correction was used and only significant differences are shown in B, C ((*) p-value < 0.05; (****) p-value < 0.0001; (n. s.) p-value >= 0.05).

A closer look into the younger and older sperm pools did not show differences in mutation frequencies between these two pools (Figure 4B); although, the mutation frequency in the younger donor pool was slightly higher than in the older donor pool (5.0 × 10^−7^ vs 4.1 × 10^−7^, respectively). However, some interesting trends can be observed in the older donor pools such as a higher mutation frequency in the Immunoglobulin-like domains I, II and III (IgI-III), but a lower frequency in the TK domain, as well as a higher frequency of substitutions with a reported germline disorder. These trends suggest that older donors might harbor more substitutions associated with a phenotype that follow a paternal age effect.

### Mutational analysis per domain

Next, we explored if the different domains of *FGFR3* are a repository of DNM. Ninety-two variants (repeated counts occurring in the same library are not included) were distributed at equal numbers across the three main domains (IgI-III n=38, TM_Inter-domain n=21, and Tyrosine Kinase n=33); yet normalized by mutation frequency, the tyrosine kinase (TK) domain had the lowest mutation frequency. Interestingly, this domain had the largest proportion of deleterious mutations (Figure 4C and Supplemental Table S1). The lower frequencies detected in the TK domain might indicate that the high deleteriousness in this domain is not tolerated in the male germline. Similarly, in a study analyzing all possible substitutions in codon p.K650 found that highly activating substitutions associated with cancer (e.g., seborrheic keratoses) were underrepresented in sperm compared to less “oncogenic” substitutions (e.g., p.K650E and p.K650T resulting in thanatophoric dysplasia type II (TDII) and familial acanthosis nigricans, respectively) (Goriely et al. 2009).

We captured in the transmembrane (TM) and surrounding regions the highest mutation frequency of DNM associated with germline disorders (Figure 4C). Also, the proportion of missense substitutions compared to synonymous substitution was highest in this domain. Interestingly, in this domain we also observed substitutions with the highest VAF (e.g., p.Y373C, p.G380R, and p.E447K); almost 50% higher in magnitude than the median of the other variants (∼4 × 10^−5^) measured in *FGFR3*. Two of those variants (p.Y373C and p.G380R) were found only in the older donor pools and have been reported to be associated with congenital disorders such as thanatophoric dysplasia type I (TDI) or ACH. Note that a higher number of COSMIC hits are also observed in this domain (Figure 3); but not in the general population (Genome Aggregation Database (gnomAD)).

Finally, our DS show that the IgI-IgIII domains accumulated a high frequency of missense variants. According to the HGMD database, the IgIII domain harbors a large number of variants associated with congenital disorders (Figure 5). However, we only detected two out of 20 of these: c.1000G>A (p.A334T) and c.749C>G (p.P250R) causing craniosynostosis and Muenke syndrome, respectively.

**Figure 5.**
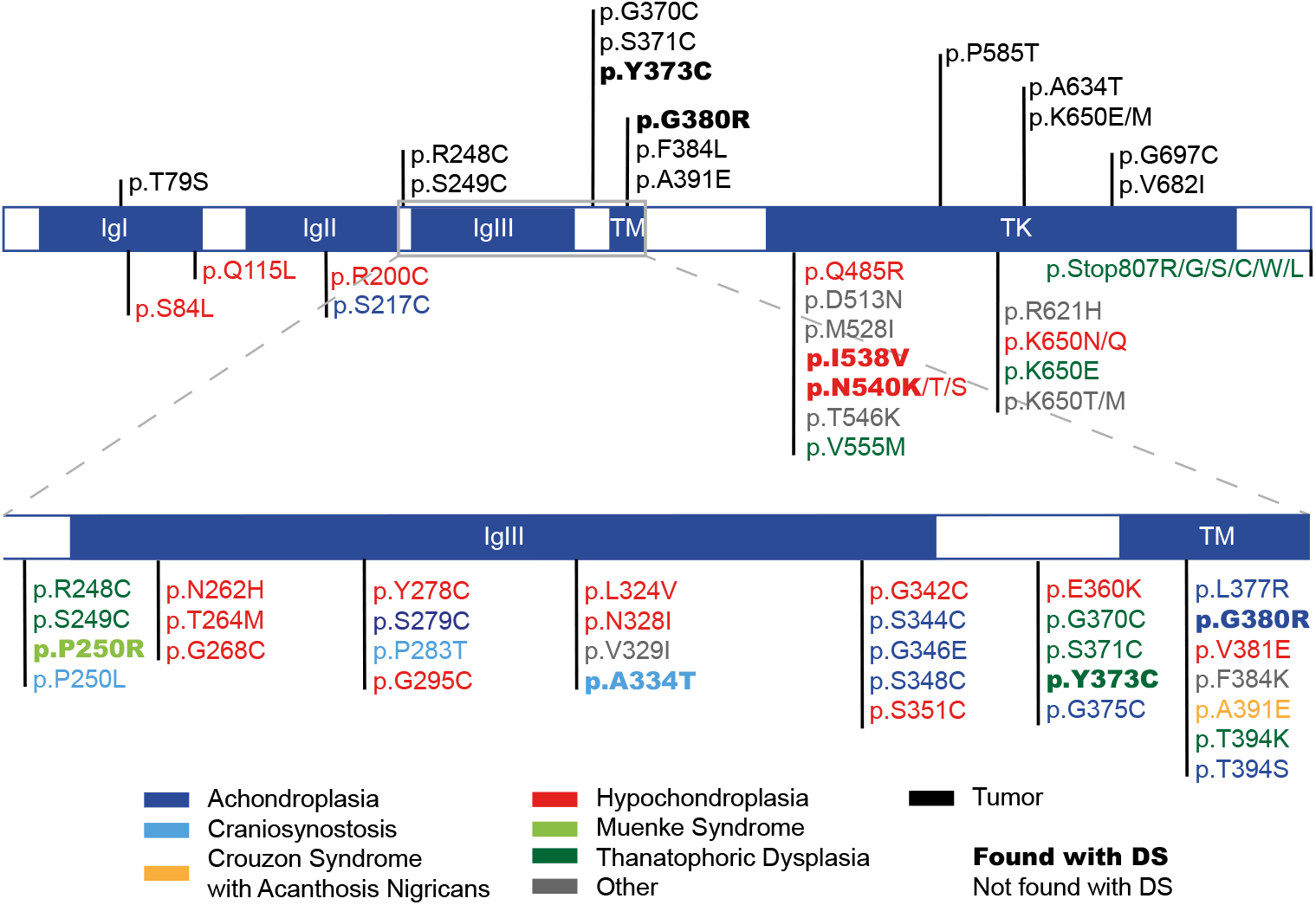
*FGFR3* variants associated with germline disorders and/or tumors. Shown are FGFR3 protein domains and associated variants (top) documented to cause germline disorders (color-coded like the respective disorder) according to the HGMD database and the variants (bottom) with the most counts in the COSMIC database. Variants captured with our DS approach are highlighted in bold. Immunoglobulin-like domain I-III (IgI-III); Transmembrane domain (TM); Tyrosine kinase domain (TK).

### Mutational mechanism

Aiming to explore the possible mutational mechanisms behind our variants, we tested if the observed variants have a selective advantage linked to the clonal expansion of driver mutations. In evolutionary genetics, the measure of dN/dS (ratio of non-synonymous versus synonymous substitutions per non-synonymous or synonymous sites, respectively) is indicative of positive, neutral or negative selection. The interpretation of this ratio is that a number close to the reference value (usually one) means no selection; a ratio below the reference means negative selection, and a ratio above the reference implies positive selection (revised in (Nielsen 2005)). The reference value is calculated by the distribution of dN/dS occurring at random with respect to nonsynonymous vs. synonymous sites (1,000 iterations). Our analysis of *FGFR3* (Supplemental Table S6) shows that there is a slight increased ratio above neutrality indicative of positive selection when taking all the data together with a dN/dS = 0.71 compared to the reference (median = 0.52), though, lying within the range of neutral expectations (lower range 0.34, upper range 0.75). The strongest indication for positive selection was in the transmembrane domain of the young donor pool with a dN/dS= 1.29 that was significantly higher to the reference (median = 0.71, lower range 0.16, upper range 1.22). Also, of interest was the IgI-IgIII domain in older donors indicating positive selection (dN/dS= 0.79, reference median = 0.48, lower range 0.24, upper range 1.06).

We further analyzed the mutational spectra, transcriptional bias, and mutational signatures in our data (Supplemental Figure S6A). Based on the cosine similarity of the *FGFR3* mutation spectra, there is a greater similarity with the COSMIC variants than with variants reported in gnomAD (cosine similarity of 0.93 vs 0.87, respectively) shown in Supplemental Table S7 and Supplemental Figure S7A. This suggests that the type of substitutions in *FGFR3* are more similar to the driver variants reported in tumors for *FGFR3* (COSMIC) than compared to random DNMs captured in the general population (gnomAD). Also, note that we do not observe differences in the mutational spectra between younger and older donor groups (Supplemental Figure S8) with both groups showing a high cosine similarity score (0.92). The lack of a strong strand bias in the substitution types also indicates that transcription cannot explain our data (Supplemental Figure S6B). Finally, the mutational signature that compares the variants also in the context of the 5′ and 3′ base adjacent to the variant shows that the main substitutions were strong to weak transitions in the context of CpG sites (VCG). Further, the comparison of our observed mutational signature with COSMIC indicate an overlap of ∼51% of the variants explained by pattern SBS24 (Aflatoxin exposure), 30% by SBS1 (spontaneous or enzymatic deamination of 5-methylcytosine to thymine) and 18.6% by SBS5 (unknown) (see Supplemental Figure S9). However, it is likely that driver genes have each their own mutational signature and needs to explored further.

Overall, the higher overall DNM mutation frequencies, the elevated VAFs at individual positions, and the trends of dN/dS for positive selection indicate that substitutions captured in *FGFR3* of sperm cannot be explained by a mutagenic mechanism similar to genome-wide DNM, but instead might be the result of deleterious/activating mutations that promote the clonal expansion of mutant cells. However, a larger dataset of variants and more sequencing data might show the underlying trends with better power.

### Testis specific clonal expansion or post-zygotic mosaic?

It is possible that the captured variants are expanding clonally with age only in the testis. Alternatively, the high VAF can also be explained by a DNM occurring during post-zygotic development thus affecting a larger number of cells (the earlier in development, the wider is the range of affected tissues), consequently originating a so-called mosaic state (Rahbari et al. 2016). Especially, mutations in genes of the RTK pathway have been documented as PZMs (reviewed in (Tiemann-Boege et al. 2020)). To distinguish between these two possibilities, we further focused on 12 variants (11 amino acid substitutions) that either (1) co-occurred in both younger and older donor pools (9 amino acid substitutions) or (2) that occurred exclusively in older donors (2 variants) at frequencies in the upper range (>1 × 10^−4^); see Table 1. The two latter variants, c.1118A>G and c.1138G>A, with a VAF of ∼3 × 10^−4^ and ∼1 × 10^−4^ respectively, were captured in multiple libraries from older donors, but not in young libraries. Interestingly, both variants are also associated with congenital disorders: variant c.1118A>G (p.Y373C) causes TDI and variant c.1138G>A (p.G380R) is linked to ACH (Figure 5). The latter has been described as a PAE-disorder with older men fathering more affected offspring ((Tiemann-Boege et al. 2002))(Shinde et al. 2013).

**Table 1.** Comparison of variants found in younger and older donor pools. Variants found in both older and younger donors are displayed together with variants found in older pools with a VAF > 1 × 10^−4^. The number of times the variant is found within a library and its corresponding DCS coverage is shown together with the average DCS coverage obtained for each variant across all libraries of the same age group. Variants with a higher frequency in older donors and variants exclusively found in older pools at a higher frequency were labelled as Paternal Age Effect (PAE) candidates; the remaining variants were considered post-zygotic mosaic (PZM) candidates. Previously described in gnomAD (1); in COSMIC (2), as congenital disorder (3), or newly discovered mutation (-); e.g., 1,3 or 2, or –. Immunoglobulin-like domain I-III (IgI-III); Transmembrane domain (TM); Tyrosine kinase domain (TK) of FGFR3.

| Younger Donors |  |  |  |  |  |  | Older Donors |  |  |  |  |  |
| --- | --- | --- | --- | --- | --- | --- | --- | --- | --- | --- | --- | --- |
| CDS | AA | FGFR3 Domain | n | DCS Coverage | Frequency | Avg DCS Coverage | n | DCS Coverage | Frequency | Avg DCS Coverage | Previously Described | Category |
| c.464G>T | p.R155L | IgII | 1 | 17840 | **5.61 x 10<sup>-5</sup>** | 13627 | 1 | 25141 | 3.98 x 10<sup>-5</sup> | 19100 | - | PZM-Candidate |
| c.841G>T | p.A281S | IgIII | 1 | 18935 | **5.28 x 10<sup>-5</sup>** | 16488 | 1 | 32262 | 3.10 x 10<sup>-5</sup> | 22784 | 2 | PZM-Candidate |
| c.952G>A | p.D318N | IgIII | 1 | 11412 | **8.76 x 10<sup>-5</sup>** | 6994 | 1 | 20014 | 5.00 x 10<sup>-5</sup> | 12857 | 1, 2 | PZM-Candidate |
| c.999C>T | p.D333D | IgIII | 1 | 7241 | **1.38 x 10<sup>-4</sup>** | 6577 | 1 | 12984 | 7.70 x 10<sup>-5</sup> | 12839 | 1 | PZM-Candidate |
| c.1118A>G | p.Y373C | Inter-domain |  |  |  | 13650 | 1 | 3169 | **3.16 x 10<sup>-4</sup>** | 8134 | 2, 3 | PAE-Candidate |
|  |  |  |  |  |  |  | 3 | 9277 | **3.23 x 10<sup>-4</sup>** |  |  |  |
| c.1138G>A | p.G380R | TM |  |  |  | 11383 | 1 | 6391 | **1.56 x 10<sup>-4</sup>** | 6586 | 2, 3 | PAE-Candidate |
|  |  |  |  |  |  |  | 1 | 8790 | **1.14 x 10<sup>-4</sup>** |  |  |  |
| c.1196G>A | p.R399H | Inter-domain | 1 | 40263 | 2.48 x 10<sup>-5</sup> | 17892 | 1 | 16741 | **5.97 x 10<sup>-5</sup>** | 12945 | 1, 2 | PAE-Candidate |
|  |  |  | 1 | 26444 | 3.78 x 10<sup>-5</sup> |  |  |  |  |  |  |  |
| c.1262G>A | p.R421Q | Inter-domain | 1 | 45305 | 2.21 x 10<sup>-5</sup> | 24745 | 1 | 19931 | **5.02 x 10<sup>-5</sup>** | 16941 | 1, 2 | PAE-Candidate |
| c.1286C>A | p.A429E | Inter-domain | 1 | 38608 | 2.59 x 10<sup>-5</sup> | 25667 | 1 | 31914 | **3.13 x 10<sup>-5</sup>** | 16573 | - | PAE-Candidate |
| c.1620C>A | p.N540K | TK | 1 | 64874 | 1.54 x 10<sup>-5</sup> | 41981 | 1 | 44819 | **2.23 x 10<sup>-5</sup>** | 27389 | 3 | PAE-Candidate |
| c.1620C>G |  |  |  |  |  |  | 2 | 28809 | **6.94 x 10<sup>-5</sup>** |  |  |  |
| c.1647G>T | p.G549G | TK | 1 | 11762 | **8.50 x 10<sup>-5</sup>** | 40410 | 1 | 17424 | 5.74 x 10<sup>-5</sup> | 27067 | 1, 3 | PZM-Candidate |
|  |  |  | 1 | 61836 | 1.62 x 10<sup>-5</sup> |  |  |  |  |  |  |  |

It is possible that variants occurring only in the older donor pool might be testis-exclusive mosaics growing larger over time (see Table 1; PAE candidates). Other interesting variants are c.1620C>A and c.1620C>G. Both are missense mutations that result in the same amino acid substitution (p.N540K) and were found 3-times in 2 different older donors libraries (VAF of 2.2 × 10^−5^ and 7.0 × 10^−5^); whereas, in younger donors libraries, they were found only once (VAF of 1.6 × 10^−5^). This amino acid substitution is known to be associated with hypochondroplasia, which also suggests that this variant could expand in the male germline with age. A more in-depth analysis of more sperm samples or the distribution within testis with a less expensive approach will provide more details in this regard.

Albeit challenging, PZM (labelled as PZM-candidates in Table 1) might be distinguished from testis-specific mosaics since they occur at a relatively high VAF already in young sperm donors or testis; whereas, PAE-candidates form no measurable clusters in testis of younger men (Goriely et al. 2003; Qin et al. 2007; Choi et al. 2012; Maher et al. 2018; Tiemann-Boege et al. 2020) and revised in (Arnheim and Calabrese 2009; Arnheim and Calabrese 2016). In Table 1, we categorized variants as PAE-candidates if detected at a higher frequency or exclusively in older donors, and as PZM-candidates if variants co-occurred in both donor pools at similar frequencies. Note that there are some variants found exclusively in younger donors. These could also be PZM-candidates missed in the older donor pool due to sampling. For those selected mutations, we further investigated if we see functional or biological differences between the PZM and PAE candidates (Table 1). Specifically, we compared the predicted deleteriousness annotated for each variant by the Combined Annotation Dependent Depletion (CADD) scores. CADD translates multiple genome annotations into a single score obtained through machine learning used as a measure of deleteriousness of single-nucleotide variants or small insertion-deletions (Rentzsch et al. 2018). As shown in Figure 6A, PAE candidates had higher CADD scores similar to COSMIC variants implying that some mutations expanding in sperm can be as deleterious as tumor-associated variants. Interestingly, PZM candidates and gnomAD variants had the lowest CADD scores. This suggests that at least part of these PAE candidate substitutions might have a stronger functional impact than most of the variants currently segregating in the population. Whether these are associated with a paternal-age effect and might have a deleterious effect like a congenital disorder, a late-onset disease (e.g., cancer), or instead a normal phenotype, is not known.

**Figure 6.**
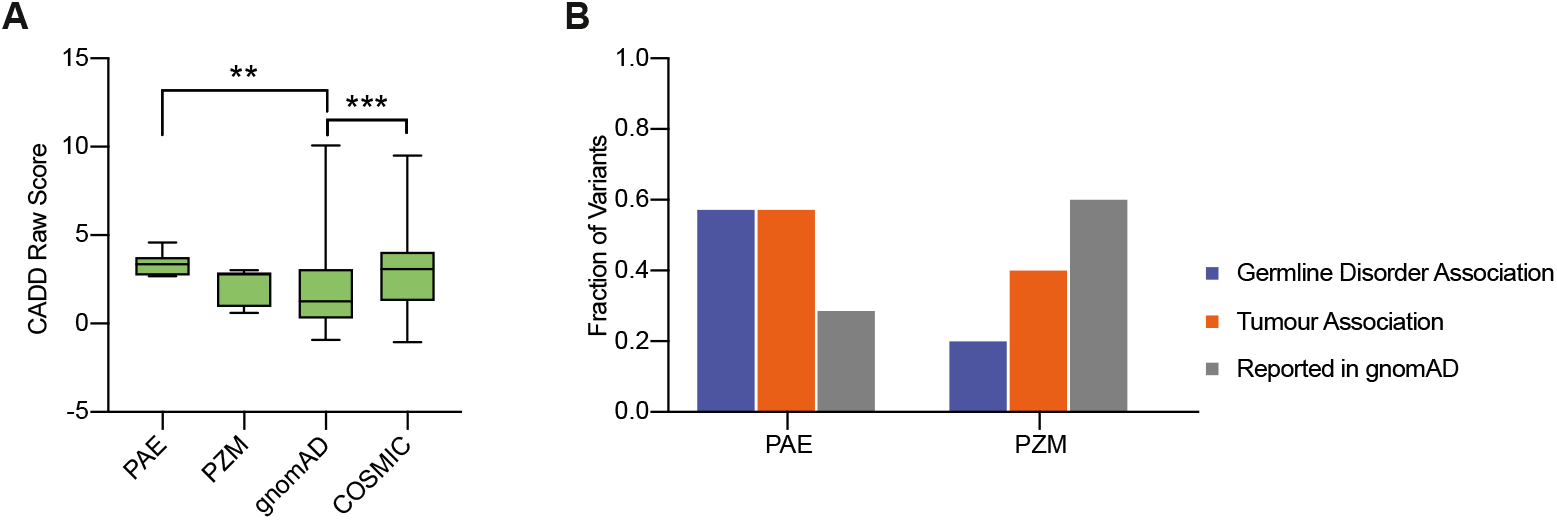
Predictive deleteriousness of variants and association with germline disorders and tumors. (*A*) Four groups of exonic single-nucleotide substitutions were analyzed: mutations selected from all sequenced libraries as potential paternal age effect (PAE, n=7); mutations selected from all sequenced libraries as potential post-zygotic mosaics (PZM, n=5); exonic single-nucleotide substitutions from gnomAD v2.1.1 database (gnomAD, n=876); exonic single-nucleotide substitutions from COSMIC v90 database (COSMIC, n=397). Boxplot comparison of the CADD raw scores obtained for each variant group. A higher score reflects a higher probability of a deleterious effect. Pairwise testing was performed using Mann-Whitney test (multiple comparison correction was applied) and only significant differences are marked ((**) p-value < 0.01; (***) p-value < 0.001). (*B*) Fraction of variants associated with germline disorders according to ClinVar; fraction of variants associated with tumors according to the COSMIC database; fraction of variants reported in the gnomAD database.

Using the same grouping of mutations, we investigated the association with germline disorders, tumors or gnomAD (Figure 5B). PAE-candidates contained the largest proportion of variants associated with germline disorders and tumors in comparison to PZM-candidates highlighting further differences between these two groups. Variants found in the general population (gnomAD) were associated more often with PZM-candidates.

## Discussion

### Targeted DS of a human gene

The establishment of DS opened exciting new possibilities in ultra-rare variant detection with an unprecedented sensitivity for a sequencing-based method (Schmitt et al. 2012; Salk et al. 2018). However, the need for consensus sequences for each DNA strand requires the sequencing of multiple PCR replicates, which translates into extremely high costs if the targeted region is large (Kennedy et al. 2014). Studies comprising the screening of mitochondrial DNA (mtDNA) have greatly benefited from the application of DS. With a length of 16,569bp, mtDNA can capitalize on the high accuracy and sensitivity of DS and still be targeted in its entirety (Kennedy et al. 2013; Ahn et al. 2015). Furthermore, in comparison with nuclear DNA, mtDNA is technically less demanding since it does not require a targeted capture strategy considering the input DNA is almost exclusively mtDNA (Kennedy et al. 2014). The most commonly used strategy to target genomic regions for DS is based on consecutive rounds of hybridization captures and enrichment (Schmitt et al. 2015; Krimmel et al. 2016; Loeb et al. 2019; Salk et al. 2019; Shen et al. 2019). Typically, the input genomic DNA is sonicated according to the chosen read length. After this step, targeted and untargeted DNA is still present. The hybridization capture strategy, although commonly used, is not efficient when dealing with small targets, with most DNA molecules being lost in the process (Winters et al. 2017; Nachmanson et al. 2018). Our DS library preparation approach used restriction enzymes to fragment DNA, followed by size selection. Like the CRISPR/Cas9 approach (Nachmanson et al. 2018), this also enables the selection of pre-determined target fragments, achieving also a low number of off-targets. Further enrichment of targeted regions is performed by two rounds of hybridization capture followed by PCR, as previously described (Schmitt et al. 2015). This approach rendered an unprecedented DCS coverage obtained for every library. Supplemental Figure S3 and S10 shows variation across different libraries, which can be explained by the different parameters used (e.g., input DNA, PCR cycles, sequencing read number). We also observed variable DCS coverage values between different targeted regions within each library that can be attributed to differences in performance of each restriction enzyme, amplification efficiency of each amplicon, and capture oligo efficiency Supplemental Figure S10. The lower DCS coverage obtained in the central parts of the restriction fragments is originated by trimming low quality read ends. However, with our DS approach we achieved a median DCS coverage of ∼11,000 allowing to call mutations at a frequency of at least 1/10,000.

### *Mutation accumulation in* FGFR3

The higher incidence of mutations in *FGFR3* increasing with age in the paternal germline was first described for achondroplasia (c.1138G>A) by screening a panel of sperm donors of different ages (Tiemann-Boege et al. 2002). Another high-resolution study that analyzed the frequency of all 9 possible substitutions in the active site of the TK domain p.K650 observed that the mutation occurrence in sperm of this particular codon is dependent on the activating effect of the variant on receptor signaling (Goriely et al. 2009). Several variants in receptor kinase receptors (e.g., *FGFR2* and *FGFR3*) expressed in the male germline have been classified as driver mutations that accumulate over time because the resulting mutated protein confers a selective advantage to the SSCs, leading to a clonal expansion of the mutant. These mutations have reported occurrences of up to 10^−4^ to 10^−5^ in sperm or testis tissue (reviewed in (Arnheim and Calabrese 2009; Goriely and Wilkie 2012; Arnheim and Calabrese 2016)).

In this study, we were able to detect 7 variants (6 amino acid substitutions) that we classified as PAE-candidates and are likely expanding in the male germline given the higher variant frequencies in older donor pools (Table 1). Three out of six of these amino acid substitutions have been associated with PAE disorders and are suspected to increase with paternal age. One of these PAE-candidates is the c.1118A>G (p.Y373C) measured at a VAF of ∼3 × 10^−4^; one of the highest frequencies measured in our dataset. This variant is causal for the autosomal dominant congenital disorder TDI, also described as a PAE-disorder (Rousseau et al. 1996b; Wilcox et al. 1998). Similarly, the c.1138G>A (p.G380R) variant measured at a frequency of ∼1 × 10^−4^ (very close to the frequency reported in other studies (Tiemann-Boege et al. 2002)) is associated with the autosomal dominant congenital disorder of achondroplasia, another PAE-disorder (Rousseau et al. 1994). This mutation is also known to increase in frequency in sperm of older donors (Tiemann-Boege et al. 2002) and it was reported to confer an advantage to the SSCs and to form clustering in the testis of an older man (Shinde et al. 2013). Another potential driver mutation rendering the amino acid substitution p.N540K was also enriched in the older donor pool. Note that for this missense mutation, we detected two different nucleotide substitutions (c.1620C>A/G). The C>A substitution was found in an older donor library with a VAF of 2.23 × 10^−5^ and the C>G substitution was detected at a VAF of 6.94 × 10^−5^. These substitutions were either lower (1.54 × 10^−5^) or absent in the younger donors pool, respectively. These two variants were also identified in a study measuring prospective mutations enriched in testis biopsies with a high RTK activity (Maher et al. 2018). The p.N540K amino acid substitution is related to the skeletal dysplasia associated with a paternal age effect, the autosomal dominant congenital disorder hypochondroplasia (Bellus et al. 1995; Rousseau et al. 1996a). The higher frequency detected in older donors coupled with an association to a PAE-disorder suggests that the p.N540K substitution might be also a testis-specific mosaic expanding with age.

Apart from the few characterized PAE-candidates, our study identified further variants that could be potentially expanding with paternal age and have not been described before. These occur mainly within or downstream of the TM domain (e.g., p.T394T, p.R399H, p. R421Q, p.A429E) and are associated with a high deleteriousness and tumors (Figure 6). Moreover, they are viable since they have been detected in the general population. Thus, our approach is an important tool to identify prospective mutations expanding with paternal age, but more evidence is required to demonstrate a selective advantage of these variants in the male germline and whether these form testis-specific mosaics with age.

### Testis-exclusive mosaics versus post-zygotic mosaicism

Driver mutations and their association with congenital disorders gain a special importance in industrial societies in which parenthood is delayed. The full understanding of driver mutations accumulating in the male germline is key to assess the risks and the impact on future generations. DNMs can occur in adult spermatogonia that are positively selected and are therefore restricted to the seminiferous tubules forming testis-exclusive mosaics. These are often associated with PAE-disorders; the numbers of mutant sperm increase with paternal age and they can be transmitted and become constitutional mutations in the offspring (Goriely et al. 2003; Qin et al. 2007; Maher et al. 2018; Tiemann-Boege et al. 2020).

We captured several substitutions with VAF over 1 × 10^−5^. These are viable variants found in the human population (gnomAD) and some of them have also been associated with tumors (COSMIC). Since they were not detected in multiple libraries and/or their frequency was not higher in older donors, we considered that these might come from a different source: DNMs occurring post-zygotically, but early in development lead to widespread tissue mosaicism where different tissues have different genotypes (Rahbari et al. 2016). A few variants detected in this study could be potential post-zygotic mutations that arose early in development, although we did not have further somatic tissue of the same donors to inspect the presence of the same variant in other tissues. Moreover, we noticed a difference in the association with germline disorders and tumors between PZM-candidates and PAE-candidates; although, our mutation deleteriousness analysis did not show any significant differences.

The expansion of PZM variants with age is not yet fully understood. In theory, the fitness of these variants conferred to the host cell should dictate their development over time. Therefore, some mosaic variants might expand clonally, and others might remain at low frequencies. But similar to gonadal-exclusive mosaics, these would also be associated with a high recurrence risk in the offspring. Depending on how early in development they occur, the range and expansion might vary (reviewed in (Tiemann-Boege et al. 2020)), but a post-zygotic mutation might be widespread and also increase with age (Salk et al. 2019) and (Acuna-Hidalgo et al. 2017). More importantly, if a DNM occurred before the separation of the somatic and gonadal lineages, then these would be equally likely to be found at high levels in younger sperm donor pools or in more pieces of testis biopsies (see such an example in (Maher et al. 2018)). Yet, in blood, the higher number of aberrant clonal expansions observed in older individuals is not fully explained by this theory. In the germline, driver mutations expand clonally with age and its frequency is higher in older individuals, but in the case of post-zygotic mutations occurring early in development, this is currently unknown. As reviewed in (Tiemann-Boege et al. 2020), different lineages of a PZM variant might have distinct fates during development. An expansion over time is observed in some tissues, but the opposite can occur in others due to a deleterious effect and subsequent decrease in relative fitness in that particular tissue.

### Effect of mutations on the RTK

To date the majority of known PAE mutations affect the RTK or components of its downstream RAS-ERK signaling pathway (FGFR3, HRAS, PTPN11, KRAS, RET, RAF1, PTPN11, BRAF, CBL, MAPK1, MAPK2) (Tiemann-Boege et al. 2002; Qin et al. 2007; Goriely et al. 2009; Choi et al. 2012; Shinde et al. 2013; Maher et al. 2016; Maher et al. 2018). Dysregulation of the FGFR3 pathway drives the preferential expansion of mutant SSCs resulting in mutant micro-clusters in testis observed with DNA-based assays (Qin et al. 2007; Choi et al. 2008; Choi et al. 2012; Shinde et al. 2013; Yoon et al. 2013) or protein markers (Goriely et al. 2003; Giannoulatou et al. 2013; Maher et al. 2016; Maher et al. 2018). The clonal expansion of PAE-mutations with age in the testis results in more mutant sperm in older individuals (Tiemann-Boege et al. 2002; Goriely et al. 2009; Maher et al. 2018), and increased incidence of the PAE-mutation.

PAE-mutations have been called “selfish” since the mutation itself, by a change in the gene product, increases its self-propagation (Goriely and Wilkie 2012). It has been postulated that the expansion of driver mutations in the testis is proportional to the hyper-reactivity of the mutant protein, with larger clusters and more mutant sperm forming with strongly activating mutations compared to mild ones (Goriely et al. 2009; Goriely and Wilkie 2012). However, the transmission of driver mutations to sperm does not always correlate with the *in vitro* activity of the RTK-RAS signaling pathway by the mutation (Giannoulatou et al. 2013), so there is no simple relationship between the activation of the mutation and the conferred strength of the selective advantage. Moreover, even though the molecular and cellular events caused by driver mutations in the male germline might parallel the events in tumorigenesis in somatic tissue, the mutational spectrum observed in tumors hardly overlaps with symptomatic spontaneous germline mutations (Aoki and Matsubara 2013; Giannoulatou et al. 2013). This difference could be explained by a tissue-specific increase in RTK-RAS signaling pathway activity. Alternatively, the mutations observed in late oncogenic stages in metastatic tissue are usually different from the initial “mild” driver mutations and change dramatically during the course of the tumor development (Sottoriva et al. 2015; Tomasetti and Vogelstein 2015). Or yet, very strong driver mutations could be detrimental in development, as it was shown for the most prevalent cancer mutation in *HRAS*, which was diminished in sperm and completely absent in birth data (Giannoulatou et al. 2013).

Accordingly, of the discovered 13 distinct variants associated with tumors, 4 might be potential PAE-candidates. Whether the remaining 9 might expand with gonadal age and contribute to viable offspring with a certain disorder or late-onset tumor remains to be tested. Interestingly, these 4 mutations are located within or nearby the TM domain of the FGFR3 protein, indicating that mutations affecting this particular domain might have a considerable impact on the protein’s activity, even in different tissues.

Not only PAE-mutations causing early-onset diseases (e.g., rasopathies or skeletal dysplasias) have been described to increase with paternal age (reviewed in (Arnheim and Calabrese 2009; Goriely et al. 2009; Goriely and Wilkie 2012; Arnheim and Calabrese 2016)). Reports have also documented an increased frequency with paternal age of late-onset disorders like breast cancer or cancers in the nervous system (Hemminki and Kyyronen 1999; Hemminki et al. 1999), or neurological and behavioral disorders, including autism (Kong et al. 2012) (Hultman et al. 2011; Frans et al. 2013), bipolar disorders (Frans et al. 2008; Grigoroiu-Serbanescu et al. 2012) or schizophrenia (Svensson et al. 2012), also reviewed in (Sharma et al. 2015; Acuna-Hidalgo et al. 2016) (Paul and Robaire 2013). Thus, the observed PAE with consequences in neurological, heart, or cancer development, might also be partially the result of DNMs in driver genes that lead to abnormalities in the signaling pathway (e.g., RTK-RAS). However, in order to better understand these effects, we need to further investigate DNMs in driver genes in the male germline, how these DNMs affect the self-propagation of the mutation, and the nature of their phenotypic consequences.

## Conclusions

With the DS approach demonstrated here we detected high frequency mutations in *FGFR3*, enabling an unprecedented view of the mutational landscape of the human male germline. Out of the 75 *FGFR3* exonic variants detected, 6 were found at higher frequency or even solely in older donors, and are distinct from the variants detected in both younger and older donors in terms of origin and phenotypic consequences. Three of them represent newly identified candidates for PAE-mutations, but further characterization is necessary. The analysis of the discovered *FGFR3* variants revealed that these mutations have a high deleteriousness compared to variants segregating in the population. Finally, our work showcases a new and accurate approach for the detection and study of DNMs in the human male germline and sheds light into different mutational mechanisms potentially affecting the receptor kinase activity and the potential risk of fathering a child at an increased age.

## Methods

### Collection of sperm and testes samples

Sperm DNA from anonymous donors mainly of Central European ancestry aged between 26 to 30 and 49-59 years were pooled using 5 donors per pool; the exact pool composition is shown in Supplemental Table S4. Samples were collected from the Kinderwunsch Klinik, MedCampus IV, Kepler Universitätsklinikum, Linz, following the protocol approved by the ethics commission of Upper Austria (Approval F1-11). Three human genomic DNA samples encoding the *FGFR3* mutations c.742C>T (NA00711), c.746C>G (NA08909), and c.749C>G (CD00002) were purchased from Coriell Cell Repositories (Camden, NJ). Similarly, genomic DNA with the *FGFR3* c.1620C>A transversion (p.540N>K) was extracted from the B-lymphocyte cell line purchased from the Coriell Cell Repositories (GM18666).

### Sample and library preparation

Sperm genomic DNA (gDNA) was extracted using the Gentra Puregene Cell kit (QIAGEN). In short, DNA of 25 μL of semen was extracted as detailed in Supplemental Materials (SM) and prepared further for the library preparation steps. DS was based on previous protocols (Kennedy et al. 2014) with substantial modifications including an initial restriction enzyme digest to pre-select the target regions. We used in the different libraries some modifications to our main protocol either starting with distinct input DNA amounts, different adapters or amplification strategies. Supplemental Table S3 summarizes the differences between each of our library preparation protocols. Genomic DNA (500-5000 ng, Supplemental Table S3) was subject to an overnight restriction enzyme digest that targeted 5 regions of the *FGFR3* gene (Supplemental Table S8). Resulting fragments are expected to be between ∼300 to ∼600 bp. A double size selection was performed using SPRIselect beads (Beckman Coulter) in order to exclude fragments outside of this size range. We used 3 different adapter synthesis protocols to produce Adapter 1, Adapter 2 and Adapter 3. The details of the protocols used to produce these adapters can be found in SM. Supplemental Figure S11 shows the structural differences between adapters: Adapter 1 is identical to the original open-hairpin structure, whereas adapters 2 and 3 have a closed loop hairpin structure that is opened up after the ligation step by the cleavage of the uracil with the USER enzyme mix (New England Biolabs). It is suggested that closed loops adapters reduce adapter dimers during ligation (New England Biolabs). Structurally, adapter 3 is similar to adapter 2, but is 9 bp longer and contains a phosphorothioate bond before the T-overhang on the 3’ end such that removal of this overhang is reduced.

Size selected genomic DNA was end-repaired and A-tailed using the NEBNext® Ultra™ II End Repair/dA-Tailing Module (New England Biolabs) according to the manufacturer’s instructions described in SM. DNA fragments with A-3’ end overhangs were then ligated to the DS adaptors with T-3’ overhangs using the NEBNext® Ultra™ II Ligation Module (New England Biolabs) following the manufacturer’s instructions and then purified by 0.8 volumes of AMPure XP beads (Beckman Coulter).

Amplification of ligated DNA was executed using KAPA HiFi HotStart ReadyMix (KAPA Biosystems). Except for 4 libraries, a strategy of 12 cycles of single primer extensions was adopted before the PCR reaction in order to get multiple copies of the initial genomic template to improve the representation of both forward and reverse SSCS and prevent the exponential amplification of templates resulting in very large family sizes. Reaction volumes and PCR conditions are described in SM and Supplemental Table S9. Primer sequences are shown in Supplemental Table S10. After the extension/PCR step, PCR products were purified with AMPure XP beads followed by 2-3 rounds of targeted capture steps to enrich further for templates of interest. A third targeted capture was done for two of the libraries using the same procedure as the second one. The number of cycles of both post-capture PCRs varied across libraries (Supplemental Table S3). Pools were verified using Fragment Analyzer HS NGS (Agilent) as described in more detail in SM.

Libraries were diluted and pooled for sequencing according to the concentration of dsDNA measured with DeNovix® dsDNA High Sensitivity. Finally, the libraries were subject to further quantification with the KAPA Library Quantification kit (KAPA Biosystems). Sequencing reactions were performed on the MiSeq Illumina platform using the kit MiSeq Reagent v3 600 cycles (Illumina) at the Center for Medical Research of the Johannes Kepler University and at the VBCF NGS Unit.

### Data processing and variant filtering

FastQ files were analyzed in Galaxy (in both public (usegalaxy.org) and private (zusie.jku.at) servers)) according to a DS specific pipeline that includes an error correction tool (Stoler et al. 2020). Variants (substitutions only) were further inspected and assigned to tiers using the Variant Analyzer (Povysil et al. 2021). Every variant was categorized as a SNP if detected in all libraries that shared the same donor pool/target region combination with a frequency ∼10%. Variants with DCS coverage below 1000, intronic variants and SNPs were discarded from our analysis and only exonic variants of Tier 1 were kept, together with Tier 2 that were detected more than once. For more details on this analysis see (Povysil et al. 2021). Selected variants were annotated using Variant Effect Predictor (VEP) (McLaren et al. 2016) and wANNOVAR (Yang and Wang 2015). The variant frequency was calculated by dividing the number of DCS calling the variant by the DCS coverage at the position of the variant within the library it was detected.

Available data on the *FGFR3* variants were extracted from gnomAD v2.1.1 (transcript ENST00000440486.2, https://gnomad.broadinstitute.org) (Karczewski et al. 2020) and COSMIC v90 (transcript ENST00000440486.7, https://cancer.sanger.ac.uk) (Tate et al. 2018) databases on January 23^rd^ 2020 and January 21^st^ 2020, respectively. Variants from gnomAD were remapped from GRCh37/hg19 to GRCh38/hg38 using NCBI Genome Remapping Service (https://www.ncbi.nlm.nih.gov/genome/tools/remap). Exonic single-nucleotide substitutions retrieved from gnomAD and COSMIC were annotated with VEP and wANNOVAR. Deleteriousness analysis was performed using CADD raw scores (Rentzsch et al. 2018) extracted from VEP annotation. Pairwise comparison between every group of variants was done with the pairwise Mann-Whitney U test and multiple comparison corrections used a False Discovery Rate (FDR) approach (two-stage set-up method of Benjamini, Krieger and Yekutieli). Germline association was investigated using ClinVar data extracted from wANNOVAR output. Tumor association was investigated by consulting the presence or absence of variants in the extracted COSMIC data.

### Mutation analysis

#### Mutation frequencies vs VAF

The VAF measures the mutant count on a single position and is estimated as the alternate allele count divided by the coverage (reference allele) on that position. The mutation frequency considers all positions of a specific region (region size) and represents the ratio of the number of different variants (variant count) per number of sequenced or analyzed nucleotides (given by the formula below). The latter is estimated as the sum of the mean coverage of a given library times the size of the sequenced/analyzed region (e.g., the region size was adjusted per domain or upstream-downstream targets). If the same variant occurred in different libraries, it was counted multiple times, otherwise it was considered only once regardless of the number of mutant counts within the same library.

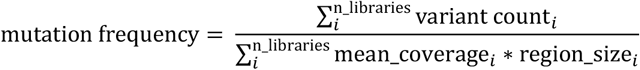

#### Mutational spectra

After counting each substitution type (e.g., A>C) we either represented the mutational spectra by the measure of relative count (variant divided by the number of all variants) or as a mutation frequency. The latter is calculated as described before, except that here we normalize for the sequenced nucleotides of the reference allele, which is estimated by the frequency of the reference allele in the targeted regions multiplied by all sequenced nucleotides.

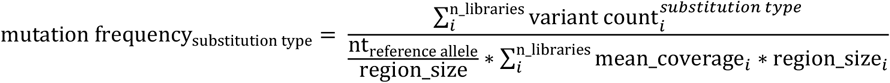

For the mutational spectra of the databases COSMIC v94 and gnomAD v3.1.1, all variants except for indels of the targeted regions were downloaded. Next, we compared them to the spectra of the duplex sequencing data using the measure of relative counts. For comparisons within duplex sequencing data (e.g., Younger and Older group), we used the measure of mutation frequencies to account also for differences in the coverage. We also categorized the mutational spectra after the (un)transcribed strand and the mutational signature within the trinucleotide context (includes the 5′ and 3′ adjacent nucleotide to the variant) with the tools SigProfilerMatrixGenerator and SigProfilerPlotting (Bergstrom et al. 2019). The mutational signature is also compared to a catalogue of signatures within the COSMIC database v3.2 (Alexandrov et al. 2020) by the tool SigProfilerExtractor (Islam et al. 2020).

#### Cosine similarities

Comparing two mutational spectra is frequently done by estimating the cosine similarity. The expected cosine similarity for a reference spectrum is calculated by bootstrapping (Abascal et al. 2021). In short, we randomly sampled *n* mutations, where *n* is the number of mutations in the query spectra, from the reference spectra and calculated the cosine similarity between the sample and the original reference spectra (1,000 iterations). The mean and 95% confidence intervals were estimated from the bootstrapped cosine similarities.

#### The rate of nonsynonymous vs. synonymous mutations

To test for selection at protein-coding sequences, we calculated the dN/dS ratio analogous to what was described in (Nielsen 2005) and (Li et al. 2015). The value of *dN* represents the number of nonsynonymous mutations per sites in the sequence where a mutation would produce a nonsynonymous difference. The value of *dS* is the number of synonymous mutations per sites in the sequence where a mutation would produce a synonymous difference. The numbers of nonsynonymous and synonymous sites was calculated using the Nei-Gojobori method (Nei and Gojobori 1986). Under neutrality, the dN/dS ratio is expected to be equal to 1, a ratio <1 is suggestive of purifying selection, a ratio >1 is suggestive of positive selection. In order to assess the probability of a random hN/hS ratios, we generated a distribution of dN/dS ratios that would be expected if the observed spectrum of mutational changes were occurring at random with respect to nonsynonymous vs. synonymous sites (1,000 iterations).

## Data Access

Raw sequencing data is available at the NCBI Sequence Read Archive (BioProject ID: PRJNA684907).

## No Competing interests

The authors declare no competing interests.

## Acknowledgements

Open access funding was provided by the Austrian Science Fund (FWF). This work was also supported by the “Austrian Science Fund” (FWFP30867000), by the FWF Doctoral College “NanoCell” (FWFW1250) and the European Regional Development Fund (REGGEN ATCZ207). Further funding was provided by the Linz Institute of Technology (LIT213201001). We would like to thank Nick Stoler, Anton Nekrutenko and Kateryna Makova for advice on duplex sequencing, and for help in the conceptualization stages of the assay.

## Authors’ contributions

I.T.-B. conceived the research; R.S., B.A, M.I, I.H., and I.T.-B. designed the experiments; R.S, S.M, T.M, A.H. and M.I performed the experiments; T.E., O.S, J.P. and I.T.-B. contributed samples, new reagents/analytic tools/data; R.S., B.A. and M.H. analyzed the data; and R.S., B.A. M.H. and I.T.-B. wrote the paper. All authors read and approved the final manuscript.

