## Supplementary Materials for "Discovery of an unusual high number of *de novo* mutations in sperm of older men using duplex sequencing"

#### Table of Contents

**Supplemental Figures** **3**

Supplemental Figure S1. Types of substitutions found either in the SSCS or the DCS. 3

Supplemental Figure S2. Age distribution of the donors used in the young and old sperm pool. 4

Supplemental Figure S3. Mutation frequencies of the sequenced libraries. 5

Supplemental Figure S4. Mutation frequencies of (non)-CpG transitions and transversions. 6

Supplemental Figure S5. Mutation frequencies of each substitution type. 7

Supplemental Figure S6. Mutational spectra (A) based on mutation frequencies, (B) transcription bias and (C) mutational signature. 8

Supplemental Figure S7. Mutational spectra of variants compared to the mutational spectra extracted from public databases. 9

Supplemental Figure S8. Mutational spectra (A) based on mutation frequencies, (B) transcription bias and (C) mutational signatures of the Younger and Older group. 10

Supplemental Figure S9. Mutational signature compared to a catalogue of signatures (version 3.2) in the COSMIC database. 11

Supplemental Figure S10. DCS Coverage of each library. 12

Supplemental Figure S11. Different adapter designs for DS. 13
Supplemental Figure S12. Example of fragment length distribution of different library preparation steps. 14
**Supplemental Methods** **16**
Collection of sperm and testes samples 16
Sample preparation 16
Duplex library preparation 17
Sequencing 22
Sequencing Data Availability 22
Data processing and variant filtering 22
Variant frequency comparison 24
Sensitivity evaluation 24
Droplet Digital PCR 25
DS vs ddPCR and BEA 25
**Supplemental References** **26**

### Supplemental Figures

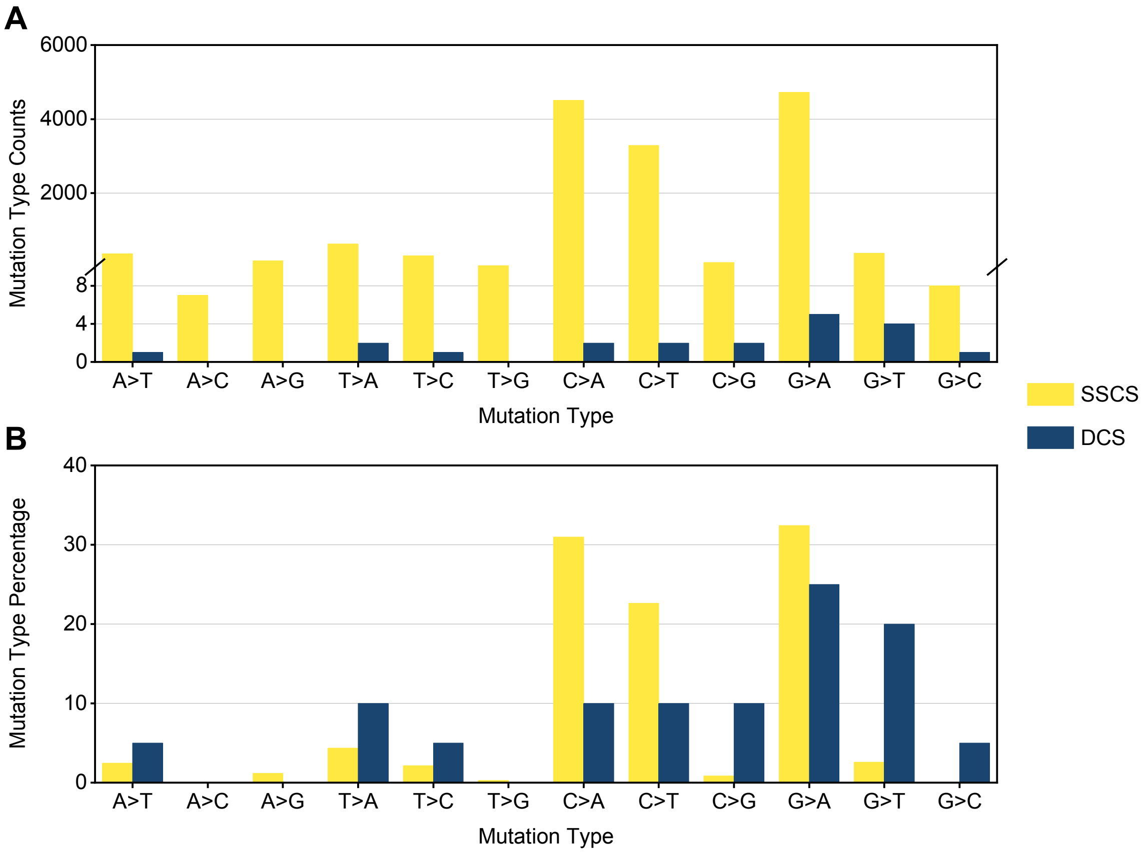


**Supplemental Figure S1.** Types of substitutions found in the SSCS and in the DCS. (*A*) Absolute number of counts of each mutation type found in libraries “FGFR3 Up O Oct19 Re-seq” and “FGFR3 Down O BAT” using SSCS (in yellow) and DCS (in blue) data. (*B*) Percentage of each mutation type found in the same libraries using SSCS data (n=14552) and DCS data (n=15).

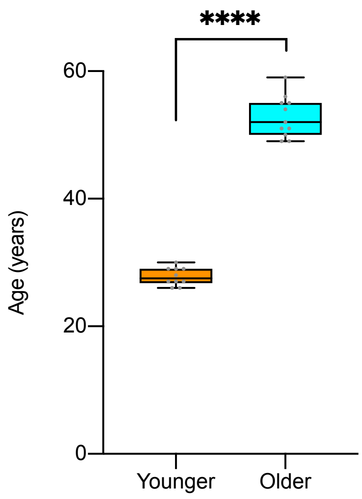


**Supplemental Figure S2.** Age distribution of the donors used in the young and old sperm pool. Each pool was composed by 5 different donors. Pools are significantly different as estimated by the Mann-Whitney U test ($P < 0.0001$).

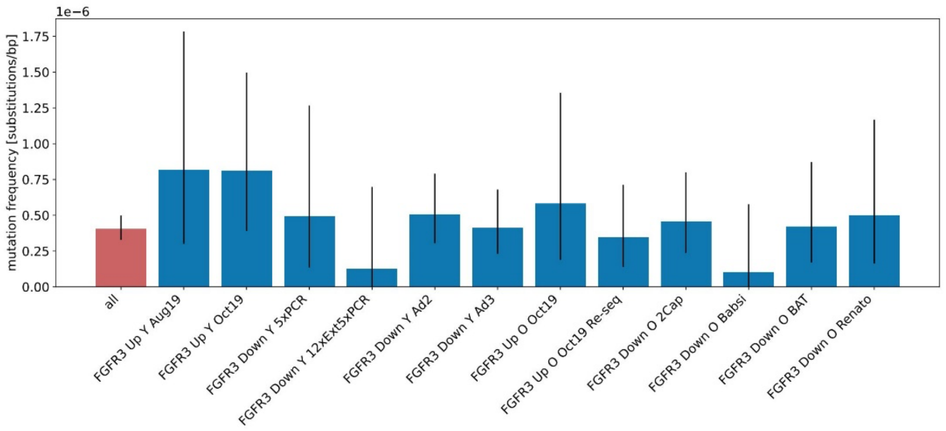


**Supplemental Figure S3**. Mutation frequencies of the different libraries estimated as the number of different variants divided by the number of total sequenced nucleotides (estimated by the mean coverage of the region multiplied with the size of the region) of that particular library. The mutation frequency of all variants (n=92) is compared to the individual libraries (FGFR3 Up Y Aug19 n=6, FGFR3 Up Y Oct19 n=10, FGFR3 Down Y 5xPCR n=4, FGFR3 Down Y 12xExt5xPCR n=1, FGFR3 Down Y Ad2 n=19, FGFR3 Down Y Ad3 n=15, FGFR3 Up O Oct19 n=5, FGFR3 Up O Oct19 Re-seq n=7, FGFR3 Down O 2Cap n=12, FGFR3 Down O Babsi n=1, FGFR3 Down O BAT n=7, FGFR3 Down O Renato n=5). Error bars are confidence intervals of a Poisson distribution. Pairwise testing between the libraries is performed with Chi-square testing with Bonferroni-Holm correction and no significance difference is found.

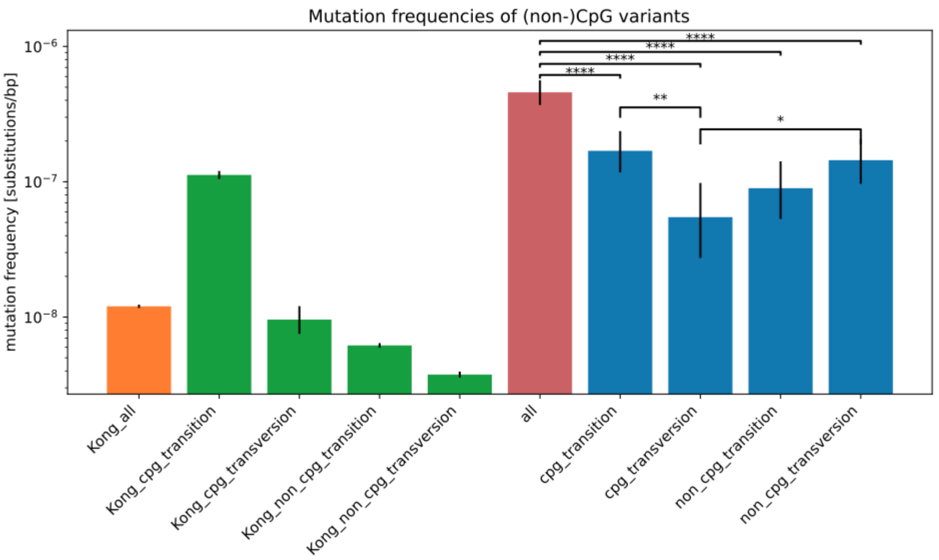


**Supplemental Figure S4.** Mutation frequencies of CpG and non-CpG variants (all n=92, CpG transitions n=34, CpG transversions n=11, non-CpG transitions n=18, non-CpG transversions n=29) compared to the pedigree sequencing of trios (Kong et al., 2012). Error bars are confidence intervals of a Poisson distribution. Pairwise testing is performed with Chi-square testing with Bonferroni-Holm correction and only significant differences are shown ((*) p-value < 0.05; (**) p-value < 0.01; (****) p-value < 0.0001).

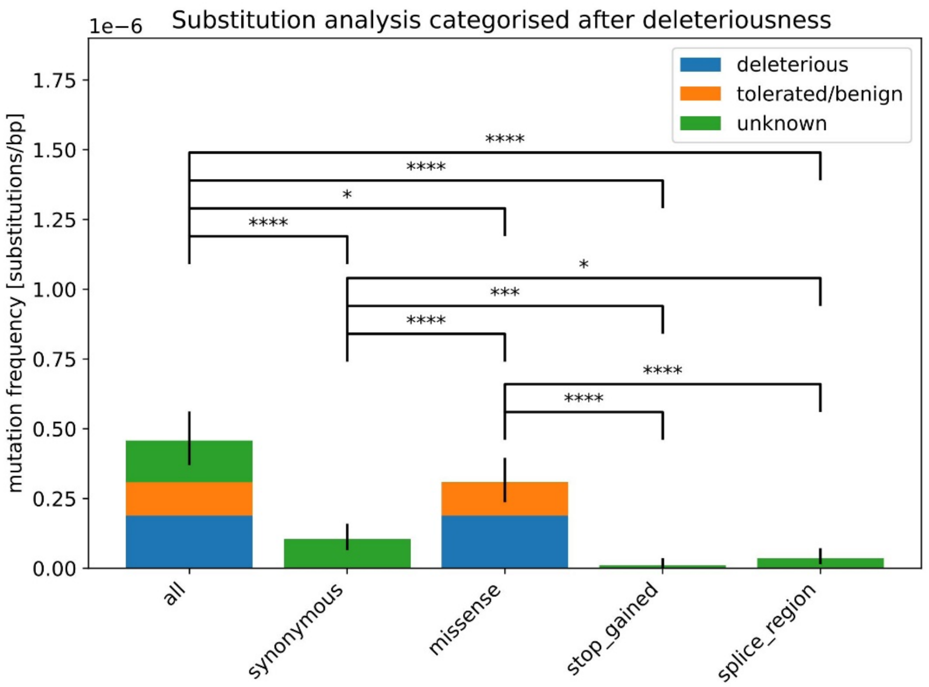


**Supplemental Figure S5.** Mutation frequencies of the different substitution types categorized after the deleteriousness based on the SIFT score (all n=92, synonymous n=21, missense n=62, stop_gained n=2, splice_region n=7). Error bars are confidence intervals of a Poisson distribution. Pairwise testing is performed with Chi-square testing with Bonferroni-Holm correction and only significant differences are shown ((*) p-value < 0.05; (**) p-value < 0.01; (***) p-value < 0.001; (****) p-value < 0.0001).

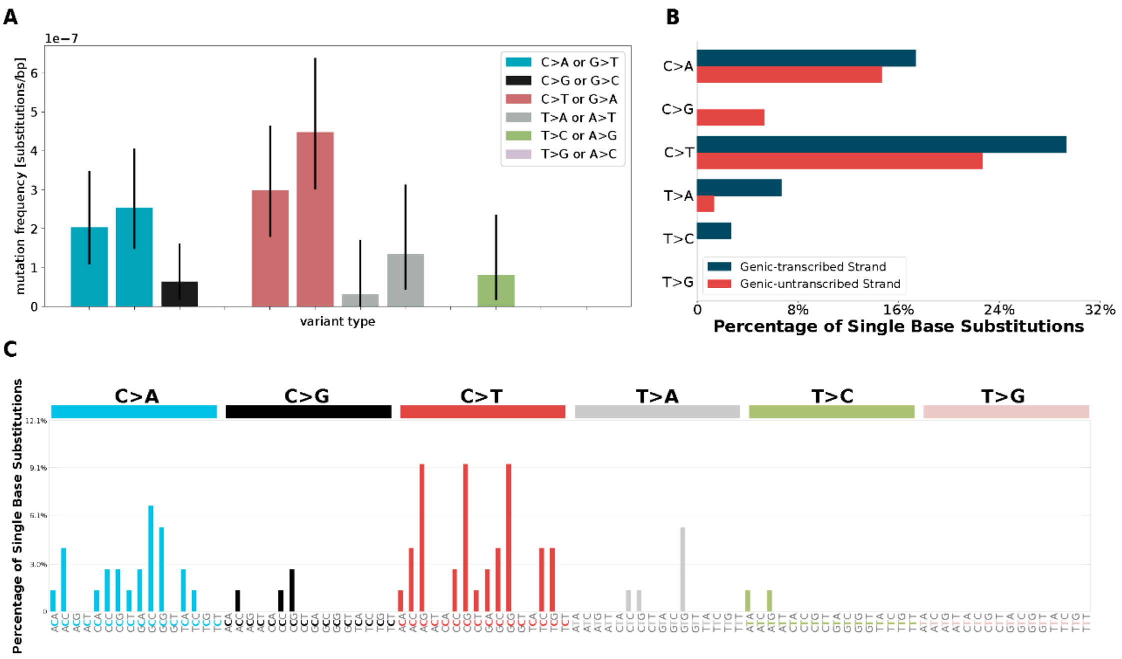


**Supplemental Figure S6**. Mutational spectra of variants (A) based on mutation frequencies (n=92) estimated as the mutation count divided by all sequenced nucleotides. (B) categorized by the transcribed and untranscribed strand. (C) Mutational signature (includes the 5' and 3' adjacent nucleotide to the variant). Note that for (B, C) every variant is considered only once (n=75) although they can be present in multiple libraries. This is required for creating the mutational spectra based on relative counts by the tools of Bergstrom et al.,2019 (Bergstrom et al. 2019).

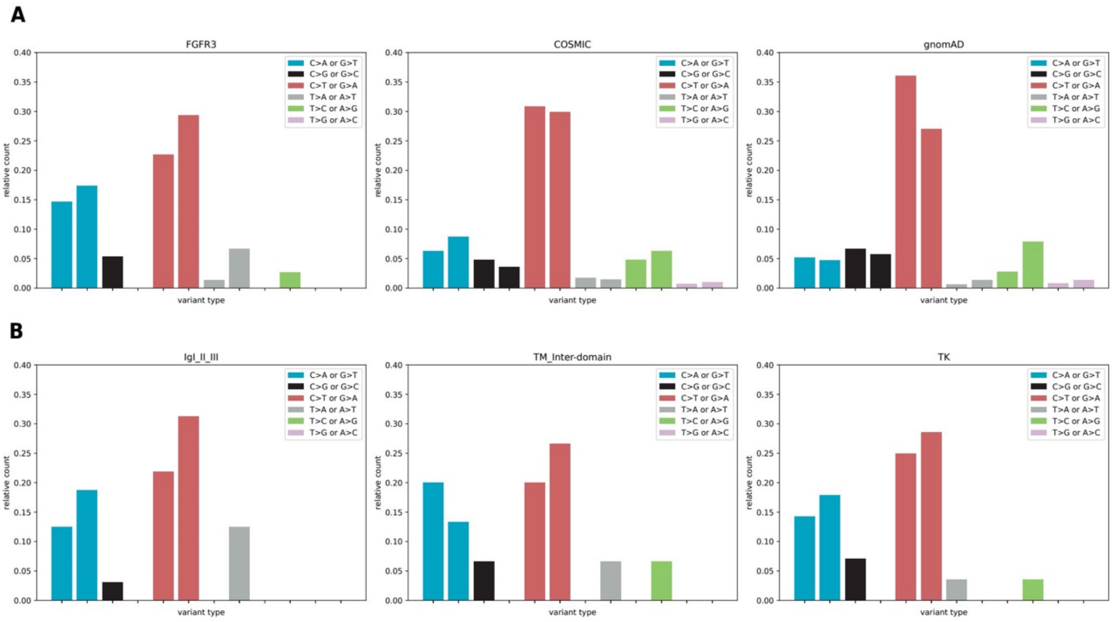


**Supplemental Figure S7.** Mutational spectra of variants compared to the mutational spectra extracted from public databases. (A) Mutational spectra of all variants (n=75) compared to COSMIC v94 (n=415) and gnomAD v3.1.1 (n=508). (B) Mutational spectra for domains (IgI-III n=32, TM_Inter-domain n=15, TK n=28). Note that the mutational spectra are based on the relative mutation count, therefore every variant is considered only once although they can be present in multiple libraires. The cosine similarities between the spectra can be found in Supplemental Table S7.

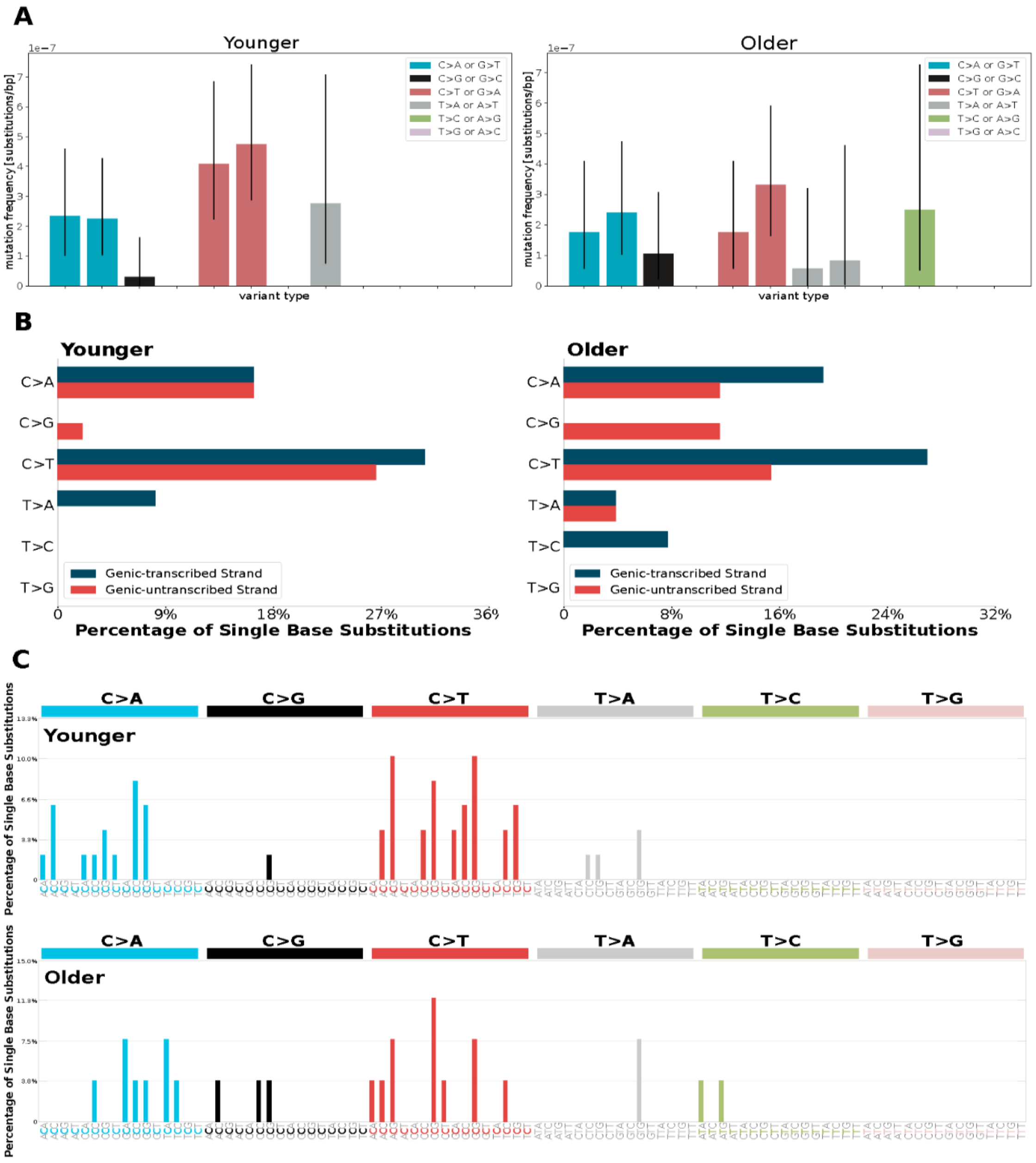


**Supplemental Figure S8.** Mutational spectra of the Younger and Older group (A) based on mutation frequencies (Younger n=55, Older n=37) estimated as the mutation count divided by all sequenced nucleotides. The cosine similarities between the groups can be found in Table S13. (B) Mutational spectra categorized by the transcribed and un-transcribed strand. (C) Mutational signatures (includes the 5' and 3' adjacent nucleotide to the variant). Note that for (B, C) every variant is considered only once (Younger n=49, Older n=26) although they can be present in multiple libraires. This is required for creating the mutational spectra based on relative counts by the tools of Bergstrom et al.,2019 ((Bergstrom et al. 2019)).

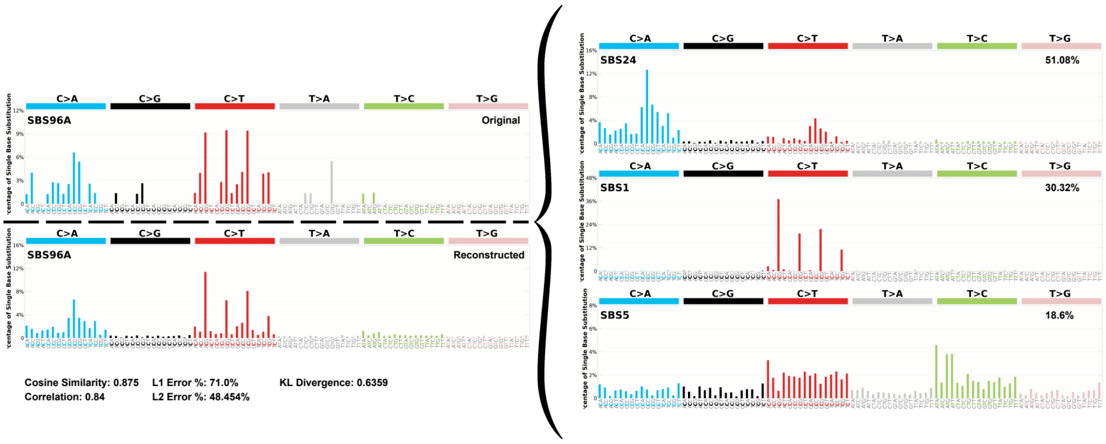


**Supplemental Figure S9.** Mutational signature compared to a catalogue of signatures (version 3.2) in the COSMIC database. The original signature can be explained by three signatures (SBS24, SBS1 and SBS5) where SBS24 (Aflatoxin exposure) contributes 51.08%, SBS1 (spontaneous or enzymatic deamination of 5-methylcytosine to thymine) 30.32% and SBS5 (unknown) 18.6% of the variants.

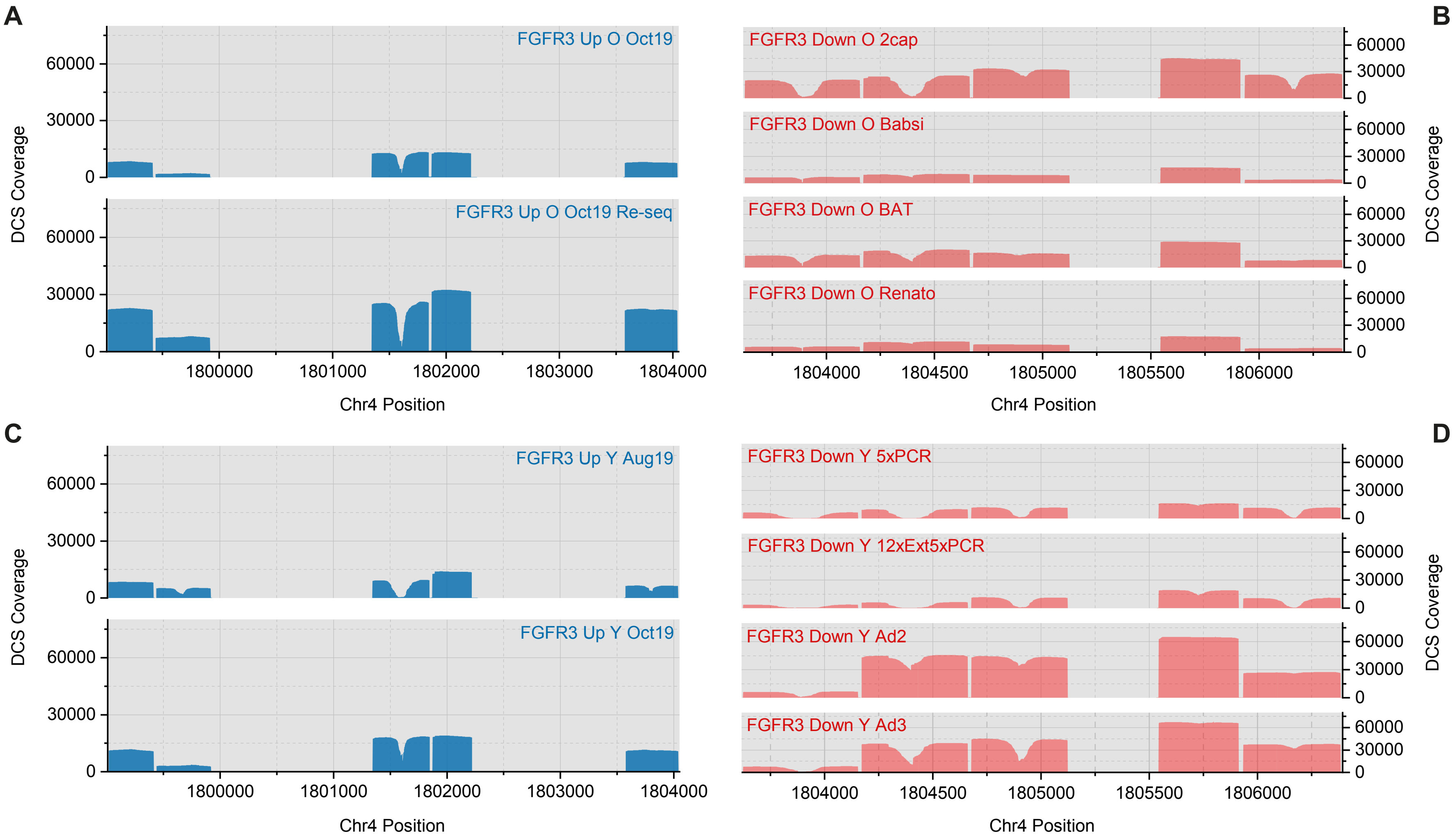
A
B
C
D
FGFR3 Up O Oct19
FGFR3 Up O Oct19 Re-seq
DCS Coverage
60000
30000
0
1800000
1801000
1802000
1803000
1804000
Chr4 Position
FGFR3 Down O 2cap
FGFR3 Down O Babsi
FGFR3 Down O BAT
FGFR3 Down O Renato
1804000
1804500
1805000
1805500
1806000
FGFR3 Up Y Aug19
FGFR3 Up Y Oct19
FGFR3 Down Y 5xPCR
FGFR3 Down Y 12xExt5xPCR
FGFR3 Down Y Ad2
FGFR3 Down Y Ad3

1

**Supplemental Figure S10.** DCS Coverage of each library. (*A*) Libraries from younger donors that targeted regions up 1-5. (*B*) Libraries from younger donors that targeted regions down 1-5. (*C*) Libraries from older donors that targeted regions up 1-5. (*D*) Libraries from older donors that targeted regions down 1-5. The variation in DCS coverage among libraries can be explained by differences in read number per library or modification in the library preparation (e.g., input DNA amount, targeted region, adaptor, hybridization capture or PCR strategy; for more details see Supplemental Methods).

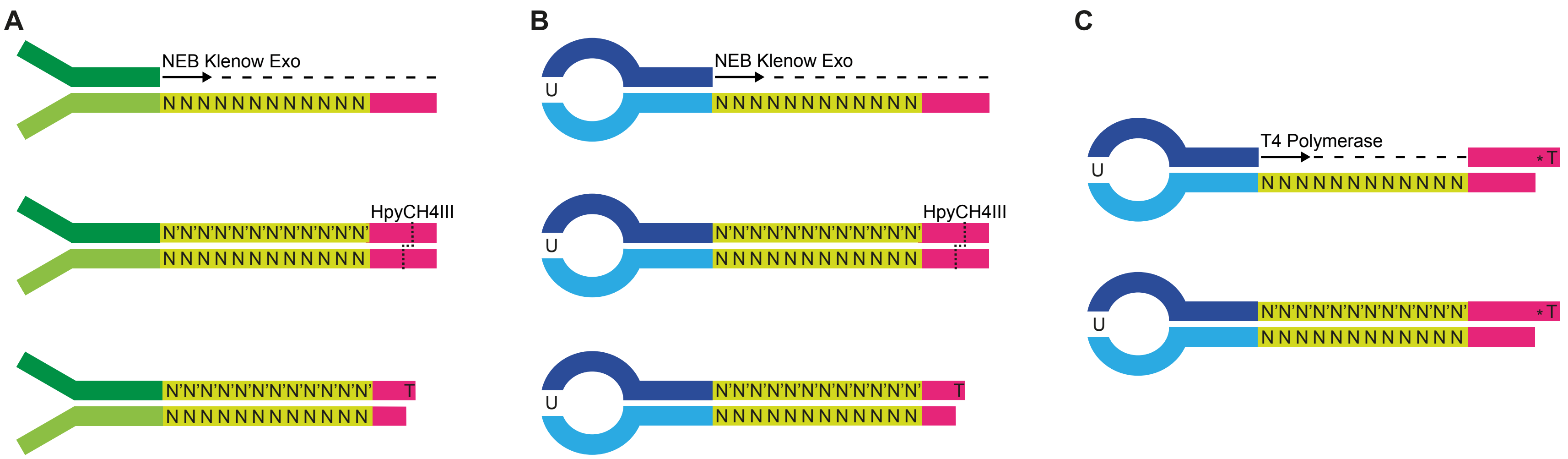


**Supplemental Figure S11**. Different adapter designs for DS. (*A*) Duplex adapter synthesis and final structure as described by Kennedy et al. (Kennedy et al. 2014). (*B*) Adapter 2 synthesis and final structure. The use of a closed loop hairpin oligo instead of the annealing of two different oligos is the only difference to the Kennedy et al. design. (*C*) Adapter 3 synthesis and final structure. This adapter uses the same looped-hairpin oligo as adapter 2 but it adds a second oligo with a phosphorothioate bond at the 3' end just before the T-overhang. After the self-annealing of the hairpin oligo and the annealing of the oligo containing the T-overhang, T4 polymerase is used to extend the hairpin oligo and to create the complement of the random barcode.

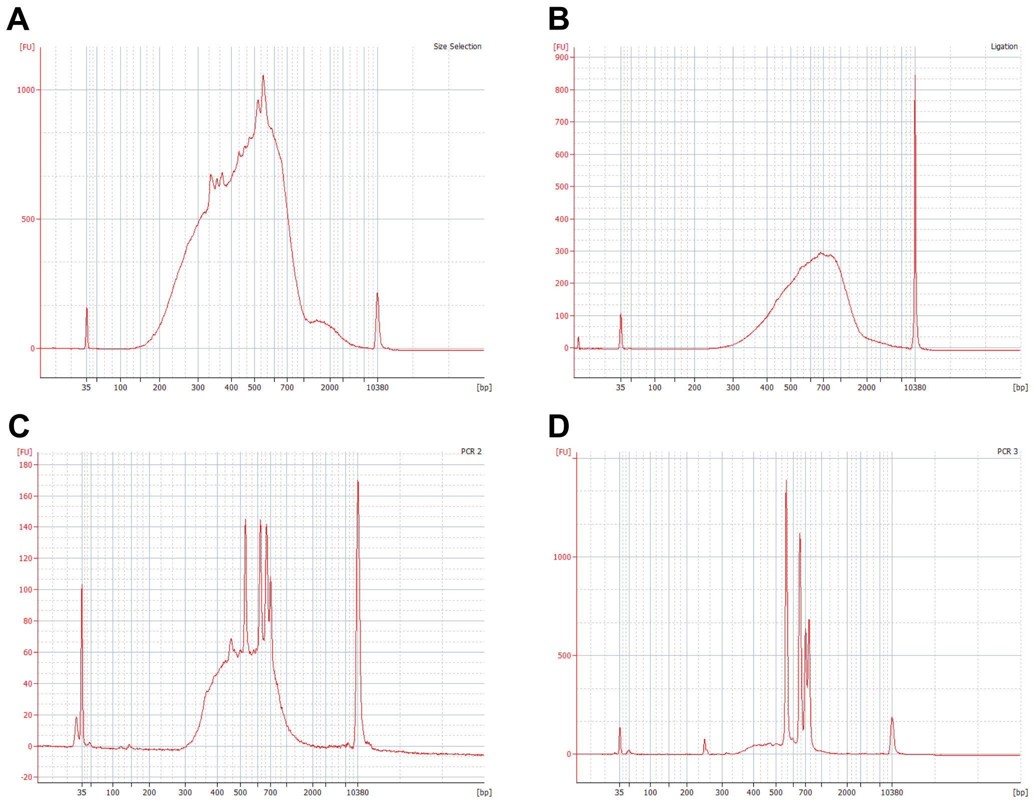


**Supplemental Figure S12.** Example of fragment length distribution of different library preparation steps. (*A*) After double size selection (~300 - 700 bp). (*B*) After ligation of adapters. (*C*) After the second PCR (performed after the first hybridization capture). (*D*) Final fragment length distribution. These examples are from library 'FGFR3 Down' O Renato and were originated by the 2100 Expert software (Agilent)

### Supplemental Methods

***Collection of sperm and testes samples***

Sperm DNA from anonymous donors mainly from Central European ancestry aged between 26 to 30 and 49 to59 years were pooled using 5 donors per pool; the exact pool composition is shown in Table S4. Samples were collected from the Kinderwunsch Klinik, MedCampus IV, Kepler Universitätsklinikum, Linz following the protocol approved by the ethics commission of Upper Austria (Approval F1-11). Three human genomic DNA samples encoding the *FGFR3* mutations c.742C>T (NA00711), c.746C>G (NA08909), and c.749C>G (CD00002) were purchased from Coriell Cell Repositories (Camden, NJ). Similarly, genomic DNA with the *FGFR3* c.1620C>A transversion (p.540N>K) was extracted from the B-lymphocyte cell line (GM18666) purchased from the Coriell Cell depository.

***Sample preparation***

Sperm genomic DNA (gDNA) was extracted using the Gentra Puregene Cell kit (QIAGEN) as follows: first, 25 µL of semen (~2 x $10^6$ sperm cells) were centrifuged (8,000g, 20 seconds) and the resulting sperm cell pellet was incubated overnight with 150 µL of cell lysis solution, 6 µL of 1M DTT, and 0.5 µL of proteinase K (20mg/mL) at 37°C. RNase treatment was performed by adding 0.75 µL of RNase A (4mg/mL) and incubating at 37°C for 15 minutes. For the protein precipitation step, the sample was put on ice for 5 minutes. Then, 50 µL of protein precipitation solution were added followed by vigorous manual shaking for 1 minute. Next, the sample was centrifuged for 20 minutes at 13,000g to remove the protein precipitate. The supernatant was transferred into a fresh tube and a second centrifugation step was performed if the solution was still cloudy. The DNA was precipitated by adding 150 µL of isopropanol (100%) and 0.25 µL of glycogen solution to the supernatant, followed by gentle inversions. A centrifugation step was done (30 minutes, 13,000g at 4°C) and the supernatant was discarded. DNA was washed with 150 µL of 70% ethanol, followed by a 3-minute centrifugation (13,000g, 4°C). Supernatant was removed and DNA was dried at room temperature for 3 minutes. Finally, DNA was resuspended in 25 µL of DNA hydration solution.

Genomic DNA from B-lymphocyte cell line was extracted using the same kit. Cells in cultured medium (~2 x $10^6$ cells) were added to a 1.5 mL microcentrifuge tube and centrifuged for 5 seconds at 13,000g. Supernatant was discarded and pellet was resuspended in residual supernatant by vortexing vigorously. Cell lysis solution was added (300 µL) followed by a 10 seconds vortexing step to lyse cells. Next, 100 µL of protein precipitation solution were added followed by 20 seconds of vortexing. The mixture was then centrifuged for 1 minute at 13,000g to pellet the proteins. Isopropanol (300 µL) was added to a clean 1.5 mL microcentrifuge tube followed by the addition of the supernatant of the previous step. The tube was inverted 50 times, followed by a centrifugation step (1 minute, 13,000g). At this stage, DNA formed a pellet and the supernatant was carefully discarded. Then, 300 µL of 70% ethanol were added and the tube was inverted several times to wash the extracted DNA. The tube was centrifuged for 1 minute at 13,000g and the supernatant was removed, followed by a drying step at room temperature for 5 minutes. Dried DNA was resuspended in 100 µL of DNA hydration solution and was vortexed for 5 seconds at medium speed. DNA was incubated overnight at 22°C with gentle shaking. Finally, samples were transferred to a storage tube.

***Duplex library preparation***

DS was based on previous protocols (Kennedy et al. 2014) with substantial modifications including an initial restriction digest to pre-select the target regions. In the different libraries we used some minor modifications to our main protocol, either starting with distinct input DNA amounts, different adapters, or amplification strategies; see Table S3. These modifications should not impact the variants called.

*Targeted enzymatic fragmentation*

Genomic DNA (500-5,000 ng, Table S3) was subject to an overnight restriction enzyme digest that targeted 5 regions of the *FGFR3* gene (Table S8). Resulting fragments were expected to have a length between ~300 to ~600 bp. A double size selection was performed using SPRIselect beads (Beckman Coulter) in order to exclude fragments outside of this size range. To the 100 µL of restriction digest solution, 93.5 µL of PCR grade water were added. Then, 106.5 µL of beads were added (0.55 volumes of beads), mixed by pipetting thoroughly and incubated at room temperature for 5 minutes. Tubes were then placed on a magnet for 5 minutes and 290 µL of supernatant were transferred to a new tube. Next, 84.2 µL of beads (1.0 volumes in total considering the initial bead solution) were added to the solution and mixed by pipetting. The mixture was incubated at room temperature for 5 minutes. Tubes were placed on a magnet and supernatant was discarded. Beads were washed twice with 80% ethanol while still on the magnet. Beads were dried at room temperature and 50 µL of PCR grade water were added. Beads were resuspended by pipetting and incubated at room temperature for 5 minutes. Finally, tubes were placed on a magnet and the clear supernatant containing the size-selected DNA was transferred to a new tube.

*Adapter synthesis*

T-tailed DS adapters were synthesized with modifications to the original protocol (Kennedy et al. 2014). We used 3 different adapter synthesis protocols to produce Adapter 1, Adapter 2, and Adapter 3. Oligonucleotides used for each adapter are listed in Table S12.

Adapter 1 was synthesized as described by (Kennedy et al. 2014), except that the amounts used for each component were downscaled. Oligonucleotides mws51 and mws55 (for each, 10 µL of 100 pmole/µL) were added to a single PCR tube and heated in a PCR machine for 5 minutes at 95°C. The PCR machine was immediately turned off and the mixture was left inside for one hour, resulting in the annealing of both oligonucleotides. The extension of annealed oligonucleotides was performed by adding 23.65 µL of PCR grade water, 5.6 µL of 10x NEB buffer 2 (New England Biolabs), 5.6 µL of 2.5mM each dNTPs (Biozym), and 1.15 µL of NEB Klenow Exo- (5U/µL) (New England Biolabs). The mixture was incubated for 1 hour at 37°C. Purification was done by adding 28 µl (0.5 volumes) of ammonium acetate (5 M) (Invitrogen) followed by the addition of 168 µL (2 volumes) of pre-cooled 100% ethanol. The solution was inverted several times until it got cloudy. Precipitation occurred while the solution was incubated for 30 minutes at -20°C. The solution was then centrifuged for 30 minutes at 4°C in a microcentrifuge at 14,000 rpm. Supernatant was then discarded and 1 mL of 80% ethanol was added followed by a 5-minute centrifugation at 4°C, 14,000 rpm. The supernatant was discarded and the tube was dried at room temperature until the pellet became transparent. The pellet was resuspended in 40 µL of PCR grade water. The T-overhang was generated by digestion with HpyCH4III enzyme: 47 µL of PCR grade water, 10 µL of 10x CutSmart buffer (New England Biolabs), and 3 µL of HpyCH4III (5U/µL) (New England Biolabs) were added to the extended adapters and incubated for 16 hours. To purify the adapters, 400 µL of PCR grade water were added, followed by 250 µL of ammonium acetate (5 M). Then, 1.5 mL of pre-cooled 100% ethanol were added and the solution was mixed by inverting until it got cloudy. Adapters were precipitated while incubated at -20°C for 30 minutes. The solution was then centrifuged for 30 minutes at 4°C, 14,000 rpm. Supernatant was discarded and 1 mL of 80% ethanol was added followed by a 5-minute centrifugation at 4°C, 14,000 rpm. The supernatant was again discarded and the tube was dried at room temperature until the pellet became transparent. Adapters were resuspended by adding 22 µL of 10mM mM $TE_{low}$ (Tris-HCl pH 8.0, 0.1 mM EDTA).

Adapter 2 was synthesized using the adapter 1 protocol. The only difference is that instead of using oligonucleotides mws51 and mws55 in the first step, only oligonucleotide DS_Hairpin_U was used, and consequently, an extra 10 µL of PCR grade water were added to the extension reaction.

The synthesis of adapter 3 started by using oligonucleotides DS_Hairpin_U and TA-Overhang. First, 10 µL of each oligonuclotide (each at 100 pmole/µL) were added to a PCR tube. The phosphorylation of 5' ends was done by adding 23.5 µL of PCR grade water, 5.2 µL of T4 DNA Ligase Buffer (10x) with ATP (10 mM) (New England Biolabs), and 3.3 µL of T4 Polynucleotide Kinase (10U/µL), followed by an incubation at 37°C for 30 minutes. Then, the solution was heated in a PCR machine at 95°C for 5 minutes. The machine was then turned off and the mixture stayed inside for 1 hour to anneal the oligonucleotides. The extension of annealed oligonucleotides was done by mixing 50 µL from the previous step with 26.7 µl of PCR grade water, 5µL of T4 DNA Ligase Buffer (10x) with ATP (10 mM), 10 µL of dNTPs (2.5 mM), 1 µL of BSA (10 mg/mL) (New England Biolabs), 0.6 µL of T4 DNA Ligase (2,000 U/µL) (New England Biolabs), and 1.7 µL of T4 DNA Polymerase (New England Biolabs). The solution was incubated at 12°C for 30 minutes followed by 16°C for another 30 minutes. The purification of adapters was accomplished by adding 400 µL of PCR grade water followed by 250 µL of ammonium acetate (5 M). Then, 1.5 mL of pre-cooled 100% ethanol were added and the solution was mixed by inverting until it got cloudy. Adapters were precipitated while incubated at -20°C for 30 minutes. The olution was then centrifuged for 30 minutes at 4°C, 14,000 rpm. The supernatant was discarded and 1 mL of 80% ethanol was added followed by a 5-minute centrifugation at 4°C, 14,000 rpm. The supernatant was again discarded and the tube was dried at room temperature until the pellet became transparent. Adapters were resuspended by adding 22 µL of 10mM mM $TE_{low}$ (Tris-HCl pH 8.0, 0.1 mM EDTA).

*End-repair/A-tailing and ligation*

End-repair/A-tailing was done by mixing 50 µL of size-selected genomic DNA with 7 µL of NEBNext Ultra II End Prep Reaction Buffer (New England Biolabs), and 3 µL of NEBNext Ultra II End Prep Enzyme Mix (New England Biolabs). The mixture was incubated at 20°C for 30 minutes followed by 65°C for 30 minutes.

The ligation of adapters occurred by adding 30 µL of NEBNext Ultra II Ligation Master Mix (New England Biolabs), 1 µL of NEBNext Ultra II Ligation Enhancer (New England Biolabs), and 2.5 µL of adapter dilution to the 60 µL of End-repair/A-tailing mix. The solution was then incubated at 20°C for 15 minutes, followed by 4°C overnight. Next, 3 µL of USER (1 U/µl) (New

England Biolabs) were added and the solution was incubated at 37°C for 15 minutes, immediately followed by AMPure XP beads (Beckman Coulter) purification. To prepare the 2.5 µL of adapter dilution, we first estimated the number of genomic DNA molecules was estimated based on the ng present after size selection, and then the amount of adapter needed to have a 20-fold excess. PCR grade water was added to fill the volume up to 2.5 µL.

Bead purification was performed as follows: 77.2 µL (0.8 volumes) of beads were added to the 96.5 µL of ligation reaction and mixed by pipetting; the mixture was incubated at room temperature for 15 minutes and then placed on a magnet for 5 minutes; the supernatant was discarded and beads were washed twice with 400 µL of 80% ethanol; the beads were dried at room temperature and resuspended by adding 20 µL of PCR grade water; the mixture was incubated at room temperature for 5 minutes and then placed on a magnet until the solution was clear; the supernatant containing ligated DNA was transferred to a new tube.

*DNA amplification and targeted capture*

The amplification of ligated DNA was done using KAPA HiFi HotStart ReadyMix (KAPA Biosystems. Components and volumes used are shown in Table S9 (PCR 1) and primer sequences are available in Table S10. Each library was split into multiple reactions, each with an input material of approximately 240 ng. With the exceptions of 4 libraries (see Table S3), the first step of amplification was 12 cycles of single primer extensions (Table S9) followed by the addition of the primer NEBNext Universal and a standard PCR amplification.

For purification, multiple PCR reactions originating from the same ligation reaction were pooled. Depending on the number of reactions and the corresponding final volume, 1.2 volumes of AMPure XP beads were mixed with each sample. The purification procedure continued as described above. DNA was eluted in 20 µL of PCR grade water.

The target capture was modified from Schmitt *et al.* (Schmitt et al. 2015). First, the purified PCR 1 output was mixed with 5 µg of Cot-I DNA (Invitrogen) and 1 nmol of blocking oligonucleotides mws60 and mws61 (Table S11). The mixture was then lyophilized and resuspended in 3.1 µL water, 8.5 µL NimbleGen 2x Hybridization Buffer (Roche) and 3.4 µL NimbleGen Hybridization Component A (Roche). Next, the mixture was heated for 10 minutes at 95°C, and 3 pmol of pooled biotinylated oligos specifically designed for the targeted regions (Table S11) were added at 65°C, followed by an incubation of 4 hours and 30 minutes. Meanwhile, 22.5 µL of NimbleGen 10x Wash Buffer I, 15 µL of 10x Wash Buffer II, 15µL of 10x Wash Buffer III, 30 µL of 10x Stringent Wash Buffer, and 150 µL of 2.5x Bead Wash Buffer

(Roche) were diluted in PCR grade water in order to obtain a 1x working solution. Wash Buffer I and Stringent Wash Buffer were incubated at 65°C. Ten minutes before the end of the hybridization reaction, streptavidin beads were washed as following: 75 µL of M-270 streptavidin beads (Life Technologies) were placed on a magnet and the supernatant was discarded; 150 µL of Bead Wash Buffer were added and the mixture was vortexed for 10 seconds and placed on a magnet; the supernatant was discarded and the wash was repeated; 75 µL of Bead Wash Buffer were added and the mixture was vortexed until homogeneous; shortly before the hybridization reaction was completed, beads were placed on a magnet, and supernatant was removed. With the beads still on the magnet, the hybridization reaction was added to the beads, quickly mixed by pipetting (it is important to be fast to maintain the temperature), transferred back to the PCR tube, and placed back in the PCR machine also at 65°C for 45 minutes. Sample were vortexed for 3 seconds every 10 minutes. After incubation, captured DNA was bound to the beads and the next step is the washing which was performed as following: with the tube still on the PCR machine, 75 µL of Wash Buffer I was added to the mixture and then vortexed for 10 seconds; the mixture was placed on a 1.5 mL tube and then placed on a magnet; the supernatant was removed and 150 µL of Stringent Wash Buffer were added and mixed by pipetting; the mixture was incubated for 5 minutes at 65°C; Stringent Wash Buffer wash was repeated; after the second incubation, the mixture was placed on a magnet and the supernatant was removed followed by the addition of 150 µL of room temperature Wash Buffer I; the mixture was vortexed for 2 minutes and placed on a magnet; 150 µL of Wash Buffer II were added and the mixture was vortexed for 1 minute and then placed on a magnet; the supernatant was removed and 150 µL of Wash Buffer III was added followed by 30 seconds of vortexing; the mixture was then placed on a magnet, the supernatant was removed and 44 µL of PCR grade water were added to resuspend the beads with the bound DNA.

The amplification of hybridized targets was done as shown in Table S9 (column PCR 2-4). PCR Product was purified using 1.2 volumes of AMPure XP beads (100 µL of PCR product and 120 µL of beads). Apart from the different initial volumes, purification was done as described before. For the final elution, 20 µL of PCR grade water was used.

For the second and third target captures, the same procedure was applied but only half of the amounts of Cot-I DNA, blocking oligos and biotinylated oligos were used (the remaining volume of the latter was replaced by PCR grade water). Also, the PCR reaction was slightly different because NEBNext® Multiplex Oligos for Illumina® (New England Biolabs) were used instead of NEB_mws20 primer in order to add indexes for sequencing multiple libraries in one sequencing run.

*Quality control*

Dilutions of 2 µL aliquots of every step of the library preparation were subject to dsDNA concentration measurements and fragment size distribution analysis to monitor the quality of the library preparation. To measure the concentration of dsDNA, the DeNovix High Sensitivity Assay (DeNovix Inc.) and a DeNovix DS-11 FX fluorometer (DeNovix Inc.) were used following the exact instructions of the manufacturer. The fragment size distribution was obtained using the High Sensitivity DNA Kit (Agilent) and a 2100 Bioanalyzer instrument (Agilent) according to the exact instructions of the manufacturer. Analysis of results was carried on with the 2100 Expert Software (Agilent). Figure S12 shows an example of fragment length distribution for a four of the different library preparation steps. The final distribution of fragment lengths is distinctive for our approach when compared with random fragmentation methods because all fragments of each targeted region have identical length.

The last step before pooling of different libraries for sequencing was a qPCR using the Library Quantification kit (KAPA Biosystems), following the procedure detailed by the manufacturer. The results were used to adjust the concentrations previously obtained with the fluorescence method. The adjusted concentrations were then used to calculate the volumes of each library that should be pooled together and sent for sequencing, following the guidelines of the sequencing provider used.

***Sequencing***

Sequencing was performed on the Illumina MiSeq platform using the MiSeq Reagent v3 600 cycles kit (Illumina) at the Center for Medical Research of the Johannes Kepler University, Linz, Austria, and at the VBCF NGS Unit, Vienna, Austria.

***Sequencing Data Availability***

Raw sequencing data is available at the NCBI Sequence Read Archive (BioProject ID: PRJNA684907).

***Data processing and variant filtering***

FastQ files were analyzed in Galaxy (on both public (usegalaxy.org) and private (zusie.jku.at) servers) to process the data according to a DS specific workflow that is available at https://usegalaxy.org/u/jku-itb-lab/w/ds-fgfr3-2021. The workflow link contains information on versions used for each tool. This workflow combines the toolset *Du Novo* (Afgan et al. 2016; Stoler et al. 2016; Stoler et al. 2020) with a recently published set of tools. These are designed not only to monitor the quality and efficiency of each library, but also to re-examine variant calls with a tier-based strategy, giving the user a better overview and control over the mutations called (Povysil et al. 2021). The first step of the workflow is grouping together all reads that share the same barcodes, which is done by the tool *Du Novo*: Make families. Next, the *Du Novo*: Correct barcodes tool will group together barcodes with differences up to 3 nucleotides (this setting can be changed) since those are probably originated by PCR or sequencing errors and would otherwise be lost (Stoler et al. 2020). The tool *Du Novo*: Align families will then perform a multiple-sequencing alignment of every read within each family of reads. The consensus sequences are then built by *Du Novo*: Make consensus reads. After trimming, consensus reads are mapped to the human genome (hg38) using Map with *BWA-MEM* (Li 2013) followed by a left realignment with *BamLeftAlign* (Garrison and Marth 2012). Variants are called using *Naive Variant Caller* (Blankenberg et al. 2014) and annotated by the Variant Annotator (Blankenberg et al. 2014). Then, the *Variant Analyzer* considers the called mutations, and re-examines the raw reads in order to assign them to different tiers based on properties such as family size or read position (Table S5). High-quality tiers (tier 1.1) require at least 3 reads per family (FS>=3); whereas, second order tiers (tier 1.2-2.4) allow also reads without a family (FS>= 1). Both assume that at least 75% of the reads in the family carry the mutant allele. Lower quality tiers (tier 3 and 4) include also other parameters such as the read position (e.g., mutants in the first or last 10bp are assigned to tier 4) or lower the percentage of reads carrying the mutant allele to >50% (tier 3).

The *Variant Analyzer* output was inspected and variants with DCS coverage below 1000, intronic variants, and SNPs were discarded. Only exonic variants of tier 1 were kept, together with tier 2 variants that were detected more than once in different libraries.

Table S14 shows an example of a partial *Variant Analyzer* output with variants found in different libraries. For this hypothetical case, we would immediately exclude variants chr4-1803735-C-A and chr4-1804278-A-T because all mutants have low quality tiers (tier 3 or 4); the high frequency and a further inspection on population databases indicates variant chr4-1804314-G-A is a SNP, so it would also be excluded; variant chr4-1803672-G-T is tier 2.1, but since it is only found once and it is not found in any other library, it would be also excluded from our data; for the remaining variants we inspect if they are exonic or intronic, and we would exclude variant chr4-1803861-G-T because it is an intronic variant for the FGFR3 transcript we are focusing on; the remaining exonic variants would then be used for downstream analysis.

Selected variants were annotated using Variant Effect Predictor (VEP) (McLaren et al. 2016) and wANNOVAR (Yang and Wang 2015).

***Variant frequency comparison***

The variant frequency was calculated by dividing the number of DCS calling the variant by the DCS coverage at the position of the variant within the library it was detected. Poisson confidence intervals (CI) were calculated for each variant according to Garwood, 1936 (Garwood 1936; Patil and Kulkarni 2012), where the lower limit is $\chi^2_{2x,\,0.025}/2$ and the upper limit is $\chi^2_{2x+1,\,0.975}/2$ and x is the number mutants detected for a particular position. CIs for each mutation frequency were obtained by dividing each of the above-mentioned limits by the number of molecules analyzed for a particular position (DCS coverage).

Available data for the *FGFR3* variants were extracted from gnomAD v2.1.1 (transcript ENST00000440486.2, https://gnomad.broadinstitute.org) (Karczewski et al. 2020) and COSMIC v90 (transcript ENST00000440486.7, https://cancer.sanger.ac.uk) (Tate et al. 2018) databases on January 23rd 2020 and January 21st 2020, respectively. Variants from gnomAD were remapped from GRCh37/hg19 to GRCh38/hg38 using NCBI Genome Remapping Service (https://www.ncbi.nlm.nih.gov/genome/tools/remap). Exonic single-nucleotide substitutions retrieved from gnomAD and COSMIC were annotated with VEP and wANNOVAR.

Deleteriousness analysis was performed using CADD raw scores (Rentzsch et al. 2018) extracted from VEP annotation.

Pairwise comparison between every group of variants was done with the Mann-Whitney test using GraphPad Prism 8.2.1 (GraphPad Software). A multiple comparison correction using a False Discovery Rate (FDR) approach (two-stage set-up method of Benjamini, Krieger and Yekutieli) was done using the same software.

Germline association was investigated using ClinVar data extracted from wANNOVAR output.

Tumor association was investigated by consulting the presence or absence of variants in the extracted COSMIC data.

***Sensitivity evaluation***

In order to assess the ability of our adapted DS approach to detect low- and ultralow-frequency variants, we sequenced four libraries with a mixture of genomic DNA containing the *FGFR3* variants c.742C>T, c.746C>G, c.749C>G, c.1620C>A, and wild-type DNA at different ratios. The ratio of each variant to WT was 1:10, 1:100, 1:1000, and 1:10,000 (one ratio per library) and it was calculated considering the heterozygosity of the genomic samples containing the variants analyzed. Variant calling and frequency calculations were done as described above.

***Droplet Digital PCR***

Site-specific assays were designed using the Droplet digital PCR Assay design tool from BioRad (available at: https://www.bio-rad.com/digital-assays). For site-specific assay information see Table S13.

Individual 20 µL ddPCR reactions were prepared by adding 10 µL of 10x SuperMix for probes (no dUTP), 2 µL of genomic DNA (~125ng/µL; total of ~72500 genomes), 6.7 µL of Nucleic Acid free water, 1 µL of primers and probes mix (900nM/probe and 100nM/primer), and 0.3 µL of restriction enzyme (CviQI or MseI; 10U/µL). Note that for each sample 4 replicates were carried out (total of ~300,000 genomes). The digest occurred at room temperature for 15 minutes prior to transferring the samples into the cartridges. 70 µL of droplet generation oil for probes and a sealing gasket were added and droplets were formed in the droplet generator. Next, the newly formed droplets were transferred to a 96-well plate and sealed for 5 seconds at 180°C. PCR was ran as follows: 10 minutes at 95°C, 40 cycles of: 30 seconds at 94°C and 1 minute at 53/55°C (depending on the target site, see Table S13), and a final step of 10 minutes at 98°C. Each step had a ramp rate of 2°C and the lid was heated to 105°C. End-point analysis was done in the droplet reader. Results were analyzed using QuantaSoft Analysis Pro Software version 1.7.4 (Bio-Rad Laboratories Inc.).

***DS vs ddPCR and BEA***

Six mutations detected with DS were measured with ddPCR or BEA in the same donors to confirm the authenticity of DS mutations. Frequencies were calculated by simply dividing the number of mutants by the number of total molecules analyzed. CIs were calculated for each variant as described above. To investigate whether there are significant differences between the results obtained with DS and ddPCR/BEA, a Fischer exact test was performed for all comparisons using the GraphPad Prism 8.2.1 software.
